# Multiparameter-based photosynthetic state transitions of single phytoplankton cells

**DOI:** 10.1101/2023.12.31.573751

**Authors:** Paul David Harris, Nadav Ben Eliezer, Nir Keren, Eitan Lerner

## Abstract

Phytoplankton are a major source of primary production. Their photosynthetic fluorescence uniquely reports on their type, physiological state and response to environmental conditions. Changes in phytoplankton photophysiology are commonly monitored by bulk fluorescence spectroscopy, where gradual changes are reported in response to different perturbations such as light intensity changes. What is the meaning of such trends in bulk parameters if their values report ensemble averages of multiple unsynchronized cells? To answer this, we developed an experimental scheme that enables acquiring multiple fluorescence parameters, from multiple excitation sources and spectral bands. This enables tracking fluorescence intensities, brightnesses and their ratios, as well as mean photon nanotimes equivalent to mean fluorescence lifetimes, one cell at a time. We monitored three different phytoplankton species during diurnal cycles and in response to an abrupt increase in light intensity. Our results show that we can define specific subpopulations of fluorescence parameters for each of the phytoplankton species and in response to varying light conditions. Importantly, we identify the cells undergo well-defined transitions between these subpopulations that characterize the different light behaviors. The approach shown in this work will be useful in the exact characterization of phytoplankton cell states and parameter signatures in response to different changes these cells experience in marine environments, which will be useful in monitoring marine-related effects of global warming.

**Significance Statement:** Using three representatives of red-linage phytoplankton we demonstrate distinct photophysiological behaviors at the single cell level. The results indicate cell wide coordination into discrete cell states. We test cell state transitions as a function of light acclimation during diurnal cycle and in response to large intensity increases, which stimulate distinct photoprotective response mechanisms. The analysis was made possible through the development of flow-based confocal detection at multiple excitation and emission wavelengths monitoring both pigment composition and photosynthetic performance. Our findings show that with enough simultaneously recorded parameters per each cell, the detection of multiple phytoplankton species at their distinct cell states is possible. This approach will be useful in examining the response of complex natural marine populations to environmental perturbations.

## Introduction

Light is the major source of energy for life on Earth. However, it is a dilute and often limiting resource for many photosynthetic organisms, including phytoplankton, arguably the most abundant source of primary productivity (1). Acclimating to light intensity alterations is therefore of principal importance for phytoplankton. Peak sunlight reaches levels of ~2.5×10^3^ mole photons, of photosynthetically active radiation m^-1^ s^-1^ (PAR 400-700nm). Nevertheless, in most aquatic environments photosynthesis takes place at much lower light intensities. Light intensity decreases exponentially, and available light wavelengths tend towards the blue end of the visible spectrum with depth in the water column. Altogether, light available in the oceans changes significantly over vertical, horizontal and temporal scales (2).

Low light intensities require efficient light harvesting and use of energy for photochemical reactions that occur within photosystems (PS). However, higher light intensities can be damaging to phytoplankton, and invoke excess energy dissipation mechanisms (3). Oceanic water columns can either be stratified, where the light intensity and quality experienced by phytoplankton cells is steady, or mixed. A vertically mixed water column results in a single phytoplankton cell being exposed to a variety of light intensities and wavelength compositions over the course of a day (4). Indeed, a diverse range of marine photoautotrophs have evolved to meet these challenges through unique light acclimation mechanisms of the photosynthetic apparatus. The form and function of the photosynthetic electron transport chain is, overall, uniform across the oxygenic photosynthetic species. Light harvesting complex (LHC) systems, on the other hand, diverge significantly between phytoplankton groups and determine the range of light intensity and quality niches they can inhabit (5).

Energy that excites photosynthetic pigments, either in the LHC or PS can be further de-excited via transfer to photochemical reaction centers (RCs) within the inner PS or dissipated. In the RCs, specific excited chlorophyll a (*chl a*) molecules undergo excited-state charge separation, initiating an electron flow that supports downstream cellular redox processes (6). Dissipation can occur in three primary ways (i) emission of a fluorescent photon, (ii) production of heat via several mechanisms collectively referred to as non-photochemical quenching (NPQ), and (iii) transition to triplet state of *chl a* which can interact with molecular oxygen to form reactive oxygen species (ROS)(7). Photosynthetic systems evolved to minimize ROS production through multiple dissipation mechanisms (8).

From a physical perspective, following excitation of a fluorophore or a pigment, the energy depletes via various pathways, whether radiatively via emitting a fluorescence photon (Rad), or non-radiatively (NR) in several ways, such as via the release of heat (H), the transfer of excitation energy to a nearby fluorophore, known as electronic excitation energy transfer (EET), through, for instance, the Förster resonance energy transfer (FRET) mechanism (9), or through the induction of a photochemical (P) reaction. Each of the above de-excitation processes per a given fluorophore has a given efficiency, also known as its quantum yield (QY), which is the ratio of the rate of these processes and the sum of all de-excitation processes (eq. 1).

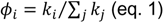

where *i* is the given de-excitation process of interest among all de-excitation processes, and *ϕ_i_* is the QY of that process. For example, the fluorescence QY of a given fluorophore can be defined as (eq. 2),

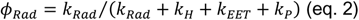

Hence, time-resolved fluorescence measurements provide direct information on the efficiencies of de-excitation processes. The typical time the fluorophore spends in the excited state until de-excitation occurs due to any of the de-excitation processes is termed the excited-state or fluorescence lifetime (eq. 3).

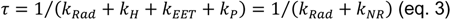

where *k_NR_* is the sum of all non-radiative de-excitation rates. Out of the above NR de-excitation processes, the rate of EET through the FRET mechanism strongly depends on the distance, *R*, from it, termed the donor, to a nearby pigment or fluorophore, which can accept the excitation energy, and thus termed the acceptor, according to (eq. 4)

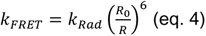

where *R*_0_ is the Förster distance, which depends on various spectral and physical parameters of the pair of donor-acceptor pigments. The FRET process, as well as other EET processes, allows excitation energy to migrate between nearby pairs of pigments until they reach the end of the cascade, populating the excited-state of RC *chl a*. Alternatively, *chl a* can also be directly excited by light. Then, *chl a* undergoes an excited-state charge transfer reaction with high QY, but can also be de-excited via the release of fluorescence or heat (eq. 5),

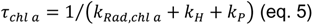

while the fluorescence lifetimes of LHC pigments prior to the RC *chl a* in the energy migration cascade is described as (eq. 6),

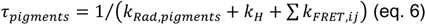

where *k_FRET,ij_* is the FRET rate for a given pair pf pigments, whether this is through the pathway of the energy migration cascade to the RC or off-pathway towards carotenoids, acting mainly as dark acceptor pigments leading to one of NPQ pathways. NPQ mechanisms are required under high light conditions to avoid the generation of ROS (7, 8, 10).

The fluorescence lifetime of directly excited RC *chl a* is well-described by eq. 5. In some wavelength ranges, additional LHC pigments, such as chlorophyll *c* (*chl c*), phycocyanin (PC), allophycocyanin (APC) and phycoerythrin (PE) have a larger cross section than *chl a*. The fluorescence lifetimes of LHC systems are well-described by eq. 6. Further, the part of their excitation energy that proceeds through the energy transfer pathway serves as a source of excitation for RC *chl a*, however with a potential delay in the excitation due to the dependence on the distance between it and the nearby pigments (eq. 4). In fact, it can be shown that the mean time spent from the moment of pigment excitation until the de-excitation of the energy from RC *chl a* depends on that pigment’s intrinsic fluorescence lifetime (i.e., the inverse of its radiative de-excitation rate, *k_Rad_*), the distance to the nearby pigment, *R*, and the RC *chl a* fluorescence lifetime (eq. 7).

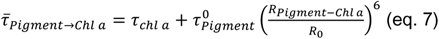

were 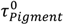 is the pigment intrinsic fluorescence lifetime. In addition, while eq. 7 considers a simplified single pair of pigment and RC *chl a*. In photosynthetic units there are ~50 PS and hundreds-to-thousands of LHC pigments, located in various positions within photosynthetic units, where each energy transfer event adds additional delay, in the form generally described by eq. 7. In addition, while many of these energy transfer events can be described as unidirectional hetero-FRET, energy transfer between same pigments, homo-FRET, can also add to the overall delay in the time spent in pigments’ excited state from the moment of excitation and until RC *chl a* de-excitation occurs (11). Nevertheless, it has been shown that any such delay in EET is expected to be negligible relative to the *chl a* fluorescence lifetime (12, 13). Notably, in FRET, the fluorescence lifetime of the pigment prior to the RC *chl a* in the migration cascade is affected by the QY of the process (eq. 8).

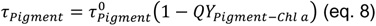

High energy transfer QY would be reflected in a short LHC pigment lifetimes, and vice versa. In general, the model developed here is a variation of the model suggested previously by Holtzwarth and co-workers, where the LHC are considered separately from the RC(6). These models suggest that effects on the QY of photochemistry will be detected in the *chl a* lifetime, and in the lifetimes of pigments prior to RC *chl a* in the energy migration cascade. In summary, fluorescence lifetimes can close “the energetic budget for the QY of photochemistry” (for a detailed discussion see Gorbunov and Falkowski (1)).

Several studies employed fluorescence lifetime approaches for the study of phytoplankton in the laboratory and in the field. These studies were performed as bulk spectroscopy, where all cells are measured together and yield ensemble-averaged results, which could be difficult to interpret (14–16). While these ensemble-averages are across many photosynthetic units in complex natural populations, a first step in better understanding the underlying distribution is by measuring these parameters one cell at a time and reporting the distribution of parameter values. Several attempts at developing single cell approaches for the study of phytoplankton primary productivity were made, including fluorescence intensity-based fast repetition rate experiments, in which important photosynthesis parameters were established (17). Additionally, there are a few examples using fluorescence lifetime imaging (FLIM) (18) in cyanobacteria and green algae (19–21), and once in a single cell flow based system (22).

As one can already understand the shear complexity of these macromolecular systems are difficult to study, even with the wealth of distinct spectroscopic features of the different pigments. Additionally, different phytoplankton species differ not only by the pigment composition but also by their amounts and ratios, and to add even more complexity these quantities are expected to dynamically vary as a function of biotic and abiotic conditions. Nevertheless, probing photosynthetic units with multiple pulsed excitations, collecting fluorescence from multiple spectral ranges and disentangling fluorescence intensity ratios from lifetimes extend the parameter space to a level in which these complexities can start being addressed.

To probe a wide range of LHC strategies, we focus on the “red lineage” oceanic phototrophs (23). Marine prokaryotic *Synechococcus* species harvest light through elaborately pigmented cytoplasmic/stromal phycobilisomes (PBSs). Their complex pigmentation pattern provides a wealth of spectroscopic handles for their function (see Fig. 1, Supplementary Table S1). Red algae are the evolutionary progeny of an endosymbiosis event with a *Synechococcus* type cyanobacteria, which also use PBSs (23). The recent resolution of *Porphyridium purpereum* red algae PBS structure (24) facilitated the ability to study their structure-function relationship (25). This massive structure contains 1,598 resolved chromophores. Dinoflagellates are secondary endosymbionts with an ancestral red alga (23). They abandoned PBS-type LHCs to utilize thylakoid-embedded *chl a/c* LHCs (apcCP) (26, 27) and lumen chlorophyll-peridinin PCP antennas (28–31) (Fig. 1, Supplementary Table S1).

**Figure 1.**
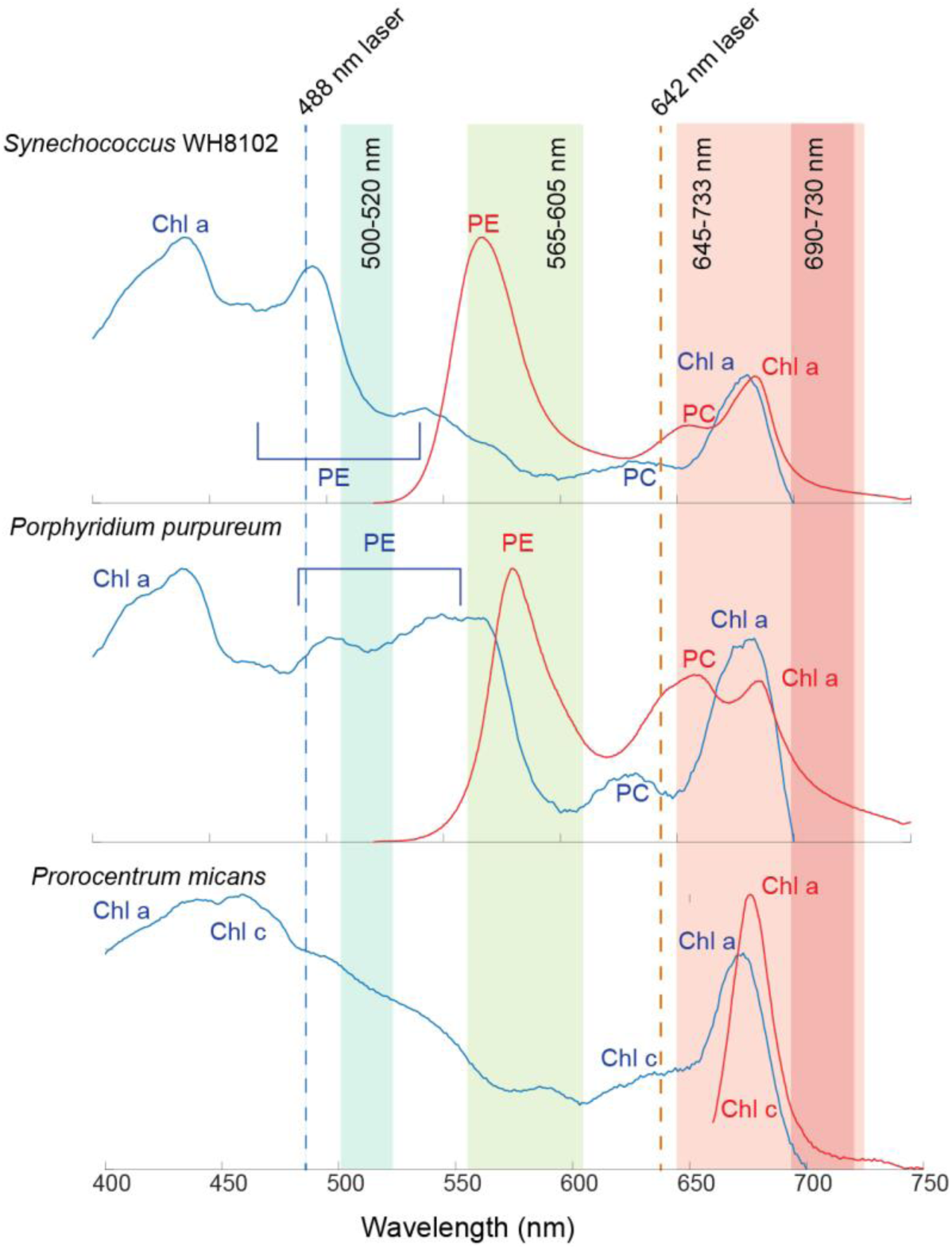
Spectra of algal species, and optical layout. The algal species used in this study: (i) *Synechococcus* WH8102 (*Syn*. WH8102), (ii) red algae *Porphyridium purpureum* (*P. purp*), and (iii) dinoflagellate *Prorocentrum micans* (*P. micans*). The absorption (blue) and the fluorescence (red) spectra are shown for each species. The spectra are normalized, and the major chromophore spectral component peaks are indicated (additional information in Supplementary Table S1). The time-resolved single cell detection system is driven by two pulsed excitation sources at 488 and 642 nm (blue and orange vertical dashed lines, respectively), and emitted photons are detected using four detectors for four separate spectral ranges (cyan, green, light red and dark red shades; additional information in tables 1, 2, and Supplementary Table S1).

**Table 1.** Selected mean fluorescence lifetime parameters.

| $\lambda_{\text{ex.}}(\text{nm})^1$ | $\lambda_{\text{detection range}}(\text{nm})^2$ | *Syn. WH8102 and P. purp* | *P. micans* |
| --- | --- | --- | --- |
| 488 | 566-605 | PE ($\tau_{488}^{PE}$) | - |
| | 645-733 | PE-excited and energy transfer to *chl a*, APC, PC ($\tau_{488}^{PC+APC+chl}$) | $\tau_{488}^{chl a+c}$ |
| | 690-730 | PE-excited and energy transfer to *chl a* ($\tau_{488}^{chl}$) | $\tau_{488}^{chl a}$ |
| 642 | 645-733 | *chl a*, APC, PC ($\tau_{642}^{PC+APC+chl}$) | $\tau_{642}^{chl a+c}$ |
| | 690-730 | *chl a* ($\tau_{642}^{chl}$) | $\tau_{642}^{chl a}$ |
| 488 & 642 | 690-730 | The mean *chl a* lifetime difference between PE-excited and energy transfer to *chl a* vs. more directly-excited *chl a* ($\tau_{488}^{chl} - \tau_{642}^{chl}$, or, $\Delta\tau^{chl}$) | - |
| <sup>1</sup> Pulsed laser excitation wavelength.<br><sup>2</sup> Fluorescence detection wavelength range. |  |  |  |

**Table 2.** Selected brightness ratio parameters.

| Calculated parameter | *Syn. WH8102 and P. purp* | *P. micans* |
| --- | --- | --- |
| $\frac{[Ex: 488, Em: (565 - 602)]}{[Ex: 488, Em: (565 - 605) \& (645 - 733)]}$ | **PE ratio**<br>PE emission brightness divided by the total emission recorded in the two channels covering emission from photosynthetic pigments. | - |
| $\frac{[Ex: 488, Em: (690 - 730)]}{[Ex: 488, Em: (565 - 605) \& (645 - 733)]}$ | **chl ratio**<br>chl a emission brightness divided by the total emissions recorded in two channels covering emission from photosynthetic pigments. | |
| $\frac{[Ex: 488, Em: (645 - 733) \& (690 - 730)]}{[Ex: 488 \& 642, Em: (645 - 733) \& (690 - 730)]}$ | **PBS cross-section (PBS C.S.)**<br>Emission brightness originating from 488 nm excitation in the chl channels divided by the sum of brightnesses resulting from excitation with both 488 and 642 nm lasers. | **Excitation cross-section (Ex C.S.)**<br>Both 488 and 642 nm excite both chl a/c. |
| $[Ex: 488, Em: (500 - 520)]$<br>(counts s<sup>-1</sup>) | **Apparent reduced Flavin content (ARFC)**<br>Some reduced flavins have emission in this spectral band. | |

To enable the direct detection of multiple meaningful single cell fluorescence parameters, we developed a multi-spectral flow-based system (Fig. 2 A). We applied this system to the study of photo-acclimation strategies in our selected model organisms: (i) a strain of the *Synechococcus* type cyanobacteria, (ii) the red algae *Porphyridium purpureum*, and (iii) the dinoflagellate *Prorocentrum micans*, in which large dynamic changes in LHC function were demonstrated (32). Our results uncover the cell-states that phytoplankton populate, their transition dynamics and efficiencies, and how they acclimate differently to light intensity changes, which were otherwise masked by ensemble averaging and the use of minimal sets of parameters. Importantly, we find that while some tested phytoplankton exhibit rapid acclimations and relaxations from light perturbation, others exhibit hours long relaxation times back from light acclimated state.

**Figure 2.**
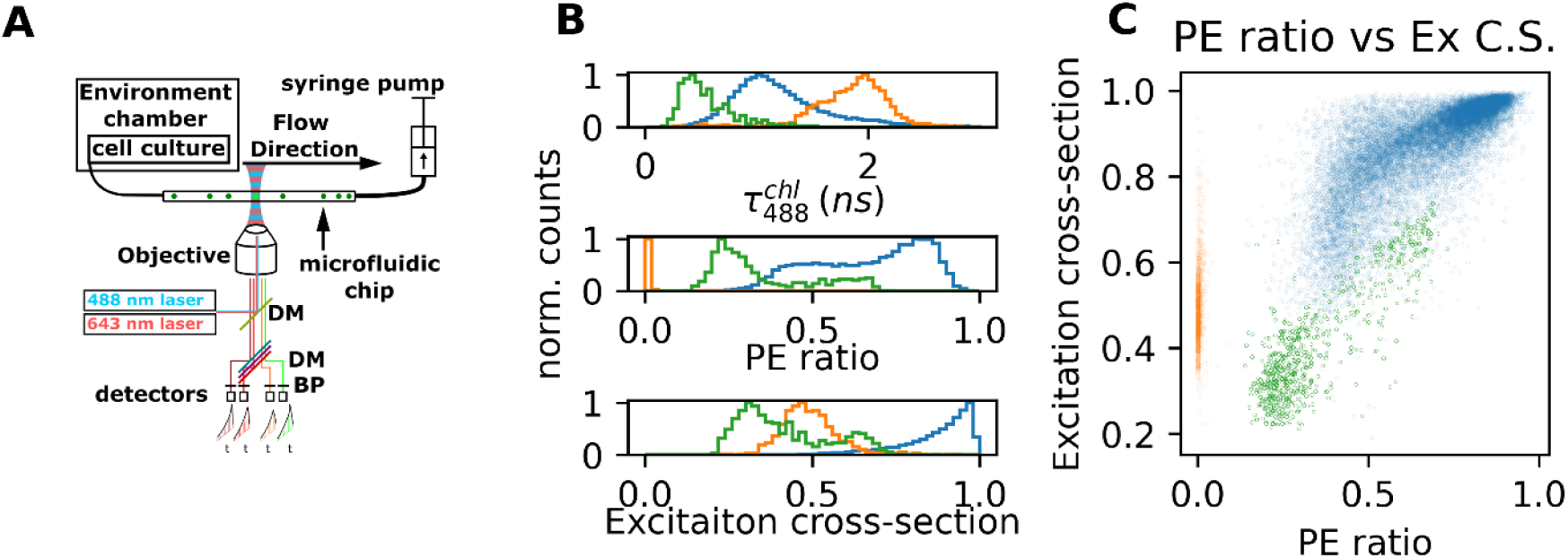
Flow-based single cell measurements. A) Diagram of the method. A cell culture is kept in an environmental chamber next to the microscope, with a culture flask containing the phytoplankton. An FPLC tube connects the culture to a microfluidic channel embedded on top of a microscope glass slide, fixed atop the microscope stage, with the outlet connected to a syringe pump, which is used to induce flow of organisms through the microfluidic channel. Pulsed-interleaved laser excitations at two wavelengths are focused into a diffraction-limited spot inside the microfluidic channel, and fluorescence photons are collected through the same objective lens, filtered, and cleaned through a pinhole. Then, the signal is chromatically split using dichroic mirrors (DM), further optically cleaned using bandpass (BP) filters and then detected in four spectral bands using single photon counting detectors. Distinct parameter signatures of different cell types are observed in B) 1D histograms of single-cell values along three selected parameters, and C) 2D scatter plot of a pair of selected parameters.

## Results

Single cell measurements were performed to uncover heterogeneity within given populations of cells, and to distinguish between different genera of phytoplankton (17). We characterized two light acclimation processes: (i) diurnal cycles, and (ii) response to an abrupt increase in light intensity.

### Synechococcus cell state transitions in diurnal cycles

Using pulsed-interleaved laser excitations at two wavelengths with four detection windows generates a large parameter space for each of the three organisms tested in this study (Figs. 1, 2, Tables 1, 2). Importantly, data collected from the different species are represented in different areas of this parameter space. This generates a clear separation of the different species in parameter space that can be easily observed in plots of selected parameters (Fig. 2, B, C).

To establish the reliability and potential of our method for reporting photophysiology at the single cell level, we choose to start by characterizing the diurnal cycle of *Synechococcus* WH8102 (*Syn.* WH8102) (Fig. 3). Daytime light intensity was set to mimic the natural light that penetrates marine water columns (60 μmol photons m^−2^ s^−1^ of blue LED light) (33), which is considered moderate for *Syn.* WH8102 (34). Small variations in mean fluorescence lifetime parameters were observed throughout the diurnal cycle (Fig. 3, A-C; Supplementary Table S2). Performing the experiment on a single-cell basis provides means of attaining cell-based parameter distributions, and hence the results are presented as parameter-based histograms as a function of time within the diurnal cycle. Examining hour-by-hour histograms of photon bursts revealed that while the distributions of mean fluorescence lifetime parameters typically showed a single clear maximum, they visually did not follow a simple Gaussian distribution, which would otherwise be expected from shot noise and central limit considerations for a single population of parameter values, suggesting at least a two-state heterogeneity of cells in the culture (Fig. 3; Supplementary Figs. S1-5). Fits to multiple Gaussian models using the F-test to determine the best-fit model with a minimal number of subpopulations similarly indicate multiple subpopulations are present (see Materials and Methods). Based on eq. 7, we can assess the QY of EET from PBS to PSII through the parameter 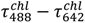 since excitation at 488 nm primarily excites PBS, while 642 more directly excites the PS *chl a*. No significant diurnal changes were detected in this parameter (Fig. 3, D), indicating that the diurnal changes observed in the individual parameters, under this light intensity, do not dramatically influence the QY of EET from PBS to PSII.

**Figure 3.**
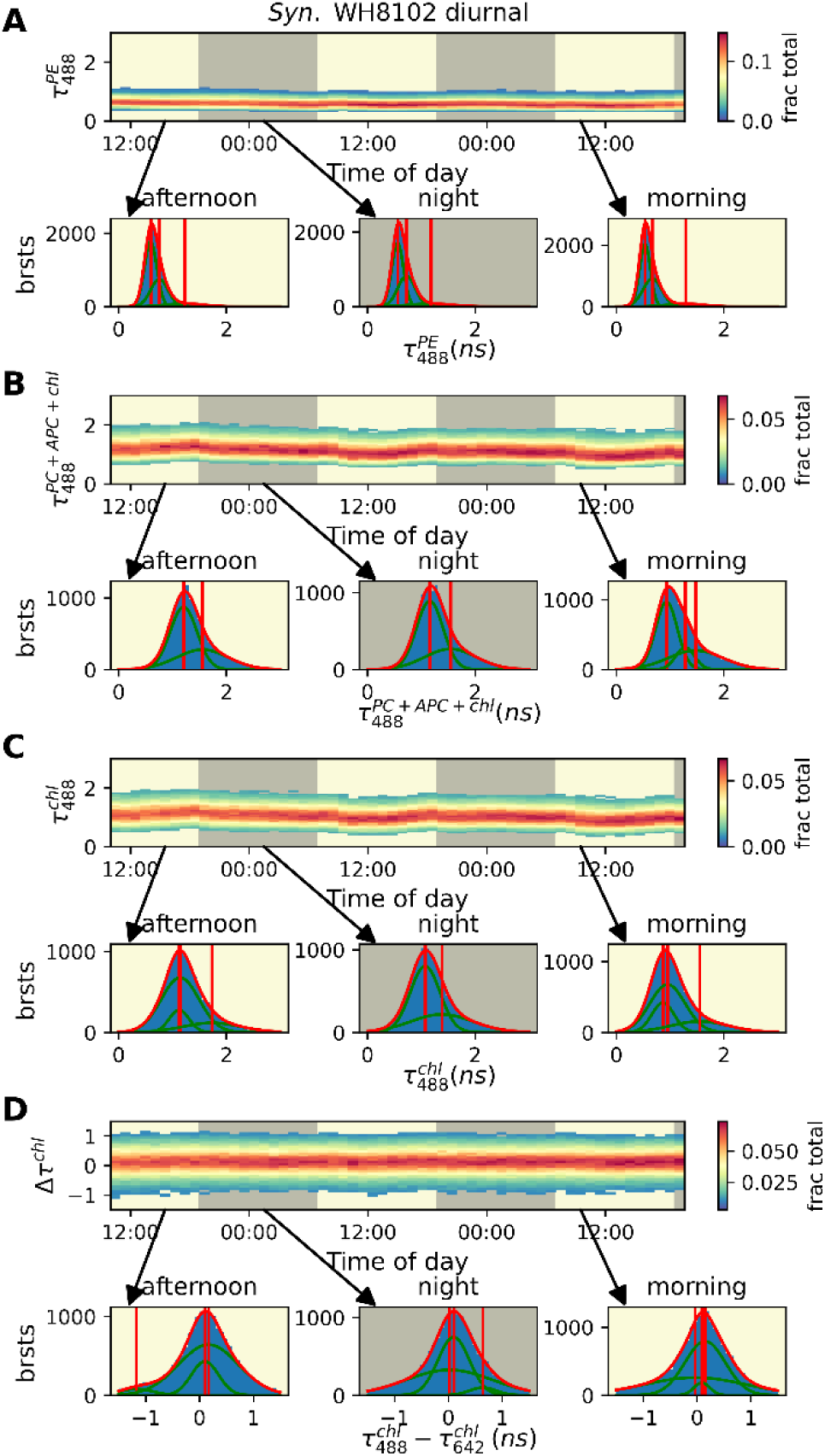
Mean fluorescence lifetime parameters in diurnal cycle of *Syn.* WH8102. Kymographs show changes in populations of mean fluorescence lifetime or lifetime difference parameters plotted along the vertical axis, and time within the diurnal cycle along the horizontal axis. Day and night are indicated by yellow and gray backgrounds, respectively. Bursts of individual organisms are grouped into hour-long bins. Plots below the kymographs show select histograms of burst mean fluorescence lifetime or lifetime difference values. Vertical lines in histograms represent the centroids of Gaussians when performing a sum-of-multiple-Gaussians model fitting to the data and selecting the best-fit model based on F-test, red line represents the fitted model, while green lines represent the Gaussians within each model. The A) mean fluorescence lifetime of 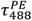 streams (i.e., direct excitation of PE), B) mean fluorescence lifetime of 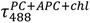 (i.e., combined excitation of PC, APC and *chl a* through PE excitation and energy transfer) C) mean fluorescence lifetime of 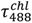 (i.e., emission of *chl a* through PE excitation and energy transfer), and D) fluorescence lifetime difference 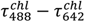 (i.e., degree of EET sensitized emission).

Next examined the spectral characteristics of the emission brightness parameter of PE (PE ratio, Table 2), where heterogeneity is found to be pronounced. Hour-by-hour histograms appear as two centrally-distributed subpopulations (Fig. 4, A-C; Supplementary Figs. S6-S8). The maxima of both subpopulations shifted slightly throughout the diurnal cycle, with more PE emission during daylight compared with nighttime. However, the shift throughout the day was smaller than the separation between the two subpopulations. When the *Syn. WH8102* PBS cross-section (Table 2) was correlated with the PE ratio, a more complex pattern emerges (Supplementary Fig. S9). There were indeed two distinct subpopulations of cells, which show two distinct PE ratio:PBS cross-section correlations. This indicates that under normal conditions the cells populate two states with distinct photophysiological behaviors. Finally, the distribution of the apparent reduced flavin content parameter (ARFC, Table 2) was found to be relatively constant throughout the diurnal cycle. The diurnal changes observed in mean fluorescence lifetime and in brightness-based parameters reflects a small degree of daytime reduction in photochemical yields that is expected under the light intensity used (34). Red algae, which use a similar yet not identical PBS LHC, showed clear diurnal oscillations in mean fluorescence lifetime parameters (Supplementary Fig. S10). In PE ratio, *chl* ratio and PBS cross-section parameters, hour by hour histograms appeared as two centrally distributed subpopulations providing clear evidence for two distinct cell states, similar to those observed in *Syn.* WH8102 (Supplementary Fig. S11).

**Figure 4.**
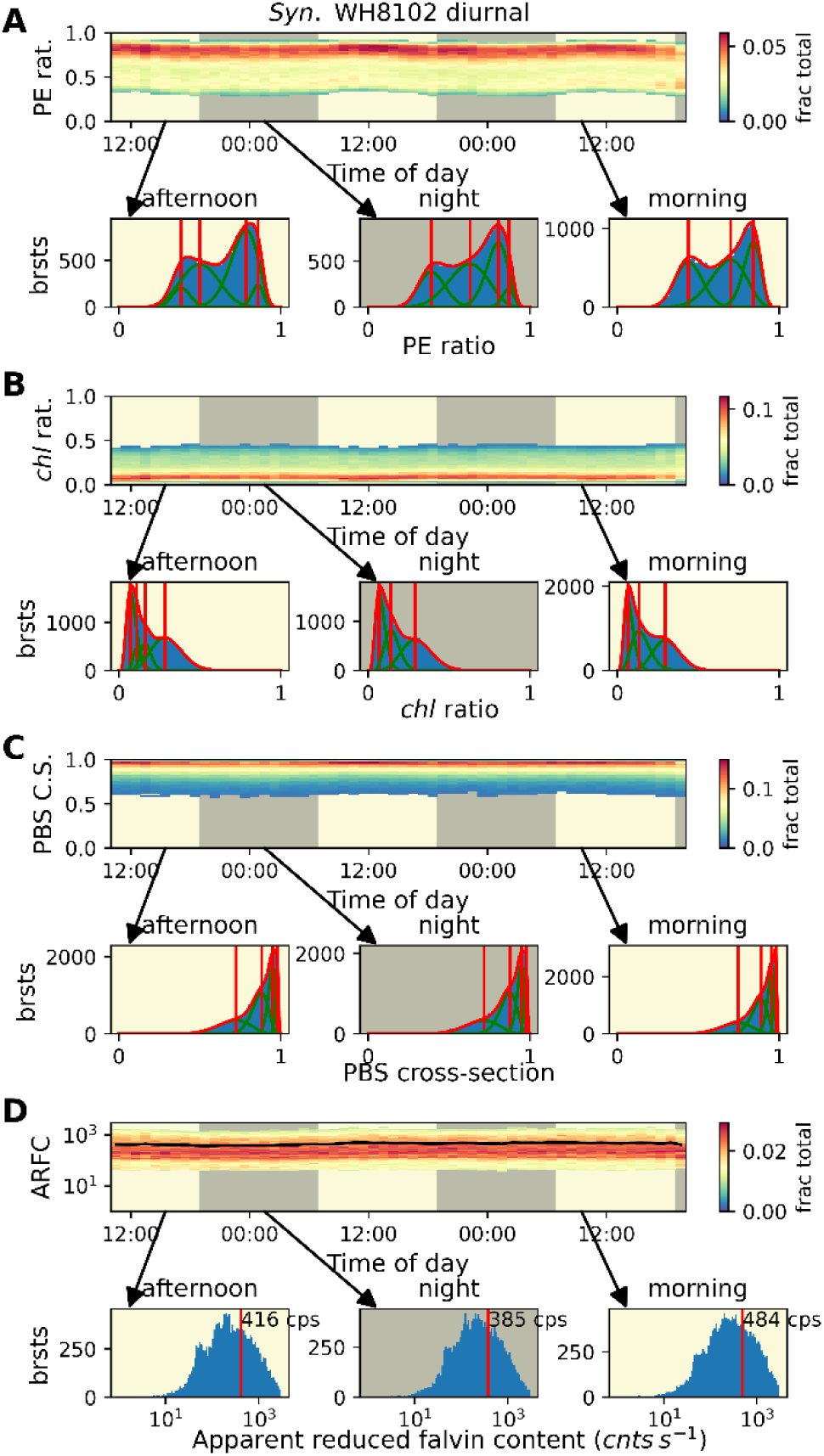
Brightness ratio parameters in diurnal cycle of *Syn.* WH8102.|. Kymographs and selected histograms of brightness and brightness ratio parameters. Displayed in the same style as Fig. 3. Bursts of individual cells are grouped into hour-long bins. Day and night are indicated by yellow and gray backgrounds respectively. Histograms of bursts below kymographs show select hour periods of brightness ratio parameters with fittings to models of a sum-of-multiple-beta-distributions, with the best-fit model selected by the F-test. Vertical lines in histograms represent the centroids of beta distributions when performing a sum-of-multiple-beta-distributions model fitting to the data and selecting the best-fit model based on F-test, red line represents the fitted model, while green lines represent the beta distributions within each model. The (A) PE ratio (relative proportion of emission from PE), (B) *chl* ratio (i.e., relative emission from *chl a*), (C) PBS cross section (relative fluorescence from 488 nm excitation), and (D) apparent reduced flavin content (i.e., brightness of emission in flavin channel), are shown. In histograms of apparent reduced flavin content the mean of raw data is reported as a vertical red line, instead of fitting the to any type of distribution.

### *Synechococcus* cell-state transitions in response to light perturbation

Next, we sought to challenge the photosynthetic apparatus by applying a light intensity perturbation. We repeated our diurnal cycle experiment, but on the second day, from 9:00 – 11:00 am, the cells were subjected to a ~5-fold increase in light intensity, which we refer to as a “light jump”. This light increase is still within the physiological range of reactions for these cells (34). Changes in parameter values were observed within the first hour of the light jump onset (Fig. 5, Supplementary Figs. S12-S20). Since the cells spend ~30 minutes in the dark while in transit from the culture flask to the confocal detection within the microfluidic channel, the response occurs within the practical time resolution of the experiment, which we substantiated using analogous bulk spectral measurements (Supplementary Fig. S21). Changes were most pronounced in mean fluorescence lifetime parameters, where a subpopulation of longer 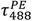 values appear with a value of ~1.6 ns (Fig. 5, A-C, Supplementary Figs. S12-S16). Two distinct subpopulations were present for all mean fluorescence lifetime parameters (Fig. 5, A-C), indicating that mean fluorescence lifetimes do not change continuously, but rather each cell belongs either to pre- or post-light jump subpopulations. Intermediate mean fluorescence lifetime values were conspicuously absent, indicating almost no cells exhibit intermediate values of this parameter. Interestingly, the long mean fluorescence lifetime subpopulation persisted for ~12 hours post light jump. Upon return to normal light intensity, the relative fraction of the long mean fluorescence lifetime subpopulation gradually decreased, yet mean fluorescence lifetime values intermediate to the two subpopulations still were not observed. After the light jump, another effect was observed in the heretofore very stable behavior of the 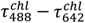 parameter. In the hours immediately following the light jump, an increase was observed in this parameter, which gradually decreased post light jump (Fig. 5, D). This suggests that much of the observed increase in mean fluorescence lifetimes was due to a conformational rearrangement of PBS and PSII, which induces a reduction of the QY of EET.

**Figure 5.**
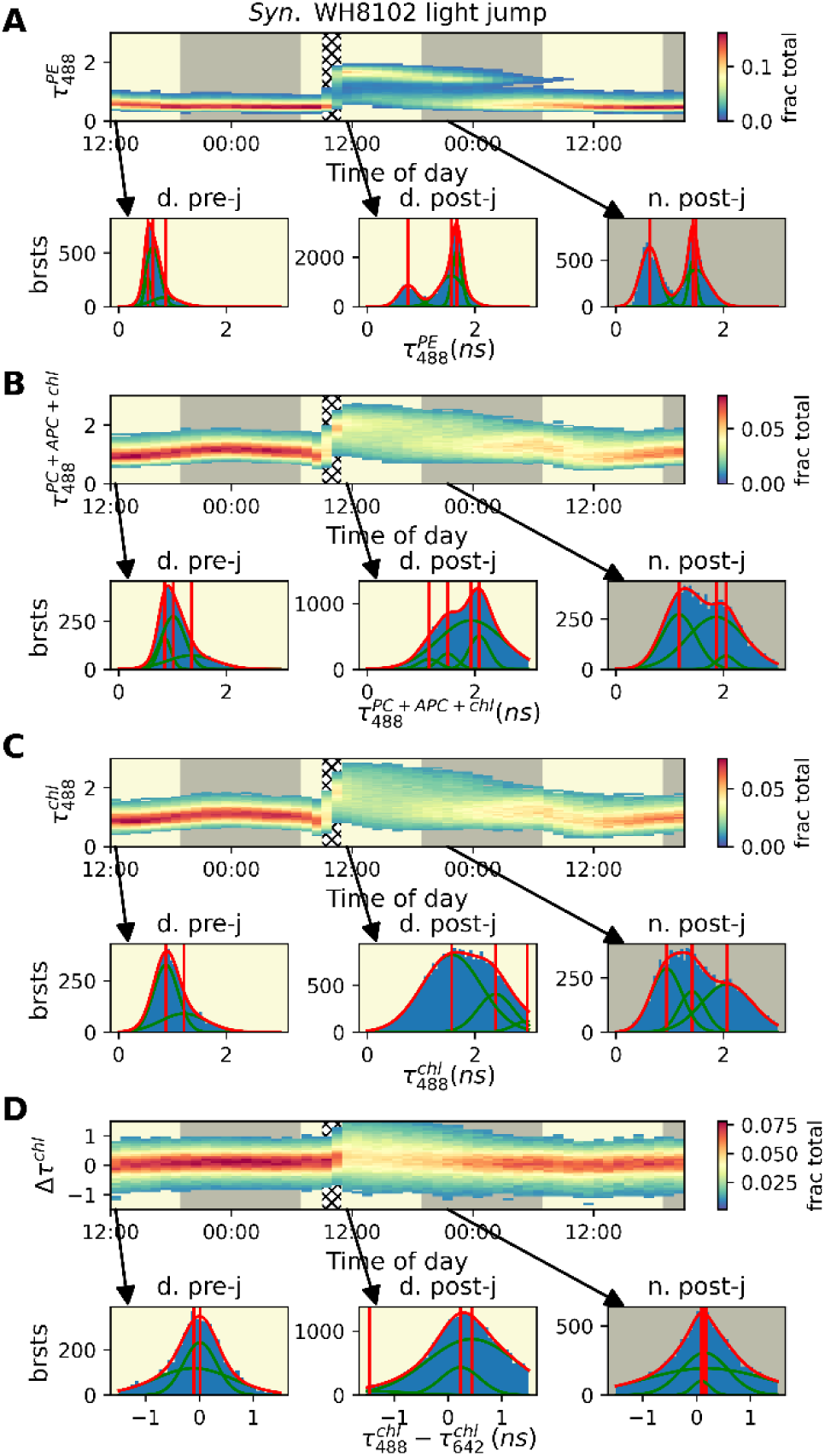
Mean fluorescence lifetime parameters of *Syn.* WH8102 in response to light perturbation. (Organized identically to Fig. 3) Kymographs show changes in populations of mean fluorescence lifetime or lifetime difference parameters plotted along the vertical axis, and time within the diurnal cycle along the horizontal axis. Day and night are indicated by yellow and gray backgrounds respectively, and the period of the light perturbation is indicated by gridded background. Bursts of individual organisms are grouped into hour-long bins. Plots below the kymographs show select histograms of burst mean fluorescence lifetime or lifetime difference values. Vertical lines in histograms represent the centroids of Gaussians when performing a sum-of-multiple-Gaussians model fitting to the data and selecting the best-fit model based on F-test, red line represents the fitted model, while green lines represent the Gaussians with each model. The A) mean fluorescence lifetime of 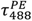 streams (i.e., direct excitation of PE), B) mean fluorescence lifetime of 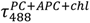 (i.e., excitation of PC, APC, and *chl* through PE excitation and energy transfer), C) mean fluorescence lifetime of 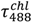 (i.e., excitation of *chl* through PE excitation and energy transfer), and D) fluorescence lifetime difference 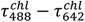 (i.e., the difference between mean fluorescence lifetimes of *chl a*, following PE excitation and energy transfer to *chl a* versus more directly excited *chl a*) are shown. This fluorescence lifetime difference reflects the rate of nonradiative energy transfer to *chl* in PSII through the PBS (PE, PC, APC).

Examination of the PE ratio histograms of the same time period showed that the subpopulation of cells exhibiting stronger *chl a* emission (i.e., lower PE ratio values; Supplementary Fig. S16) decreased in amplitude. The centroid of the high PE ratio subpopulation increased slightly during the light jump, while that of the lower PE ratio showed a more pronounced increase in its centroid, from a mean of ~0.4 during daylight before the light jump, to a mean of ~0.6 after the light jump, respectively (Supplementary Fig. S21, A, B). Bulk fluorescence measurements exhibited a small yet observable decrease in the intensity of the corresponding peaks (Supplementary Fig. S20).

By examining the PE ratio histograms filtered by 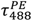 values (Supplementary Fig. S22), it is clear that the long 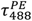 subpopulation after the light jump originated from transitions of cells from both basal short 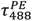 subpopulations. Additionally, the apparent reduced flavin content rose due to the light jump, and slowly returns to pre-light jump values by the beginning of the next day (Supplementary Fig. S17, D).

### The intracellular meaning of correlated state transitions

Our flow-based data establishes switching between distinct cell-states observed through multiple fluorescence-based parameters. However, the burst-based parameter values may serve as ensemble averages of different photosynthetic complexes within each cell. To test this, we further verified the above flow-based single-cell observations by cell imaging. We perform pulsed-interleaved excitation (PIE (35); also referred to as nanosecond alternating laser excitation, nsALEX (36)) coupled with FLIM, PIE-FLIM (37), acquisitions of cyanobacteria spatially-confined in agarose (Fig. 6). Performing PIE-FLIM allows us to probe the fluorescence parameters we use in the flow-based measurement, also in imaging mode. We tested if pixel-wise parameters inside the cells also distribute into distinct groups of fluorescence values. Our results indicate that Syn. WH8102 cells group into two distinct predominant subpopulations for many of our fluorescence-based parameters. Additionally, we used the white light of the microscope, typically used for recording bright field images, as our source for a light jump. In these “light jump” experiments, we are able to show the increase in mean fluorescence lifetime parameter values 30 min after the light jump event, as well as smaller changes in intensity ratio parameters and in the fluorescence lifetime difference parameter. However, the absolute values of the parameters were different from the ones observed in flow. We attribute these differences to the different conditions the cells were exposed to for establishing their spatial confinement. Conditions in the flow experiment culture flask were closer to the environmental conditions than those in agarose, where gas diffusion is limited. Imaging, therefore, qualitatively corroborates the two-state behavior of *Syn.* WH8102 cells observed in flow-based measurements. Most importantly, the cell-averaged parameter values indeed represent spatially distinct cell states.

**Figure 6.**
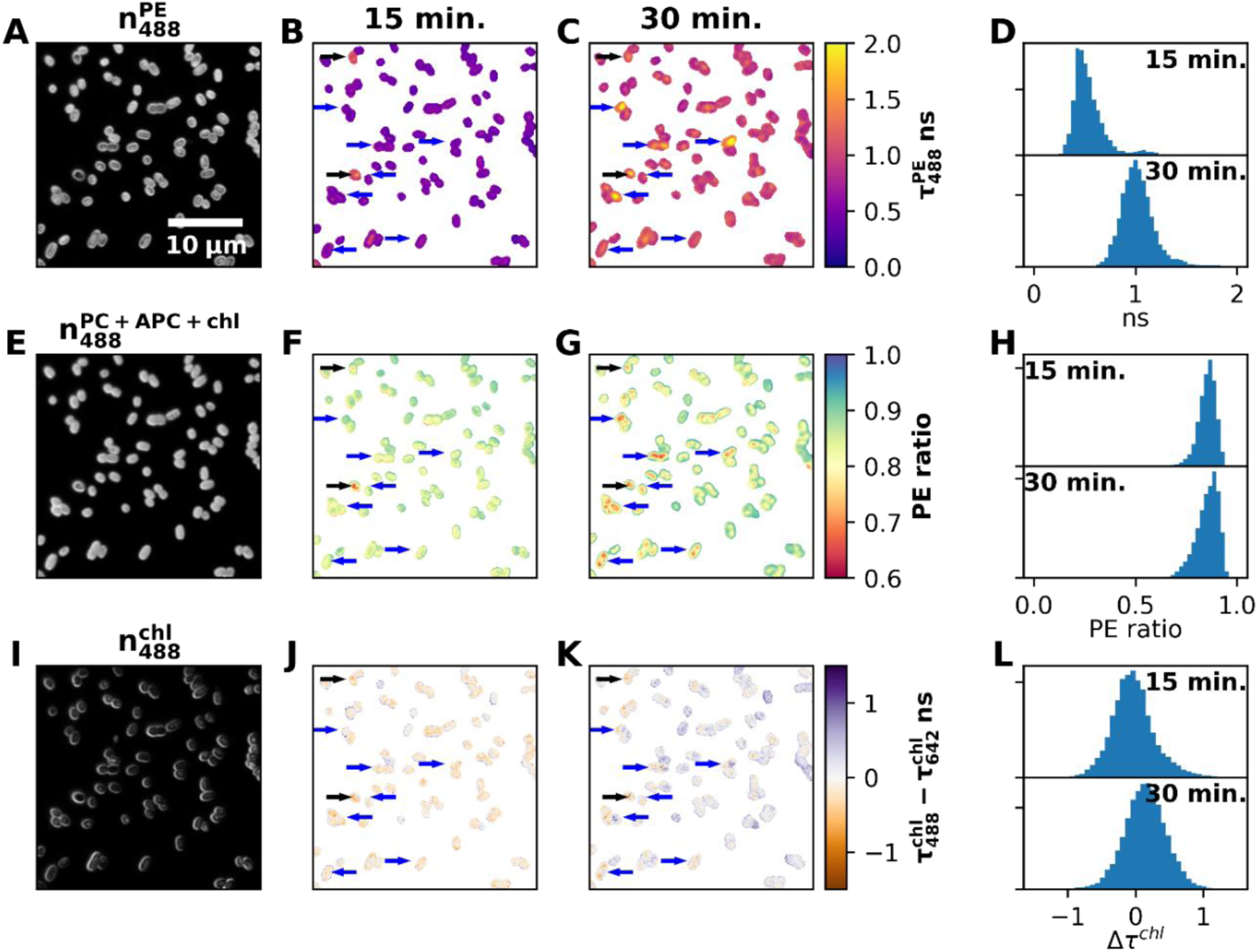
Spatial distribution of parameter values in imaging of *Syn.* WH8102. Images show parameter values in Syn. WH8102 cells for 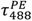 (B, C) PE ratio (F, G), and 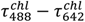 (J, K), 15 (B, F, J) and 30 (C, G, K) minutes into the light jump. Shown are also pixel-wise histograms of these parameter values (D, H, L). Values reported for cells with a threshold of 200 photons in the λ_ex._=488 nm, PC+APC+chl channel per pixel. Intensities in 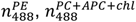, and 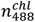 channels are shown in A, E, and I respectively. Black arrows indicate selected cells with long 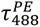 lifetimes in the 15 minute light jump condition, while blue arrows indicate selected cells with low PE ratio in the 30 minute light jump condition.

The data observed here, on a single cell basis, exhibited a similar trend to the previously-reported bulk spectroscopic parameters of light jump experiments (38). Previous experiments, however, could not have resolved the distinct fluorescence-based cell-state transitions that we report here, due to ensemble averaging over multiple unsynchronized cells.

### Cell state transitions in dinoflagellates

So far, we focused on phytoplankton with PBS systems. Seeking to extend our understanding to chlorophyll-based light harvesting systems, we turned our attention to the dinoflagellate *P. micans*, which uses c*hl a/c* LHCs. This requires different terminology for derived parameters (Tables 1,2, Supplementary Table S1). *P. micans* displayed only minor changes in the mean fluorescence lifetime parameter 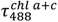 throughout the diurnal cycle with 6 μmol PAR (Fig. 7, A), and negligible in QY of EET in 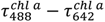 (Fig. 7 B). Owing to its almost exclusive emission in the far red, most parameters based on brightnesses and their ratios, such as the excitation cross-section, showed little fluctuations (Fig. 7, C, G). *P. micans* responded to light jumps with a decrease in 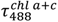 indicative of nonradiative dissipative processes, such as NPQ, while showing no change in 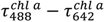 indicating no modulation of QY of EET (Fig. 7, F). Lifetime values recovered within 1-2 hours post light jump (Fig. 7, E). The apparent reduced flavin content, which remained constant throughout the diurnal cycle, was similarly perturbed during the light jump into the hour afterwards, decreasing significantly before returning to the steady-state values observed before the light jump (Fig. 7, D, H).

**Figure 7.**
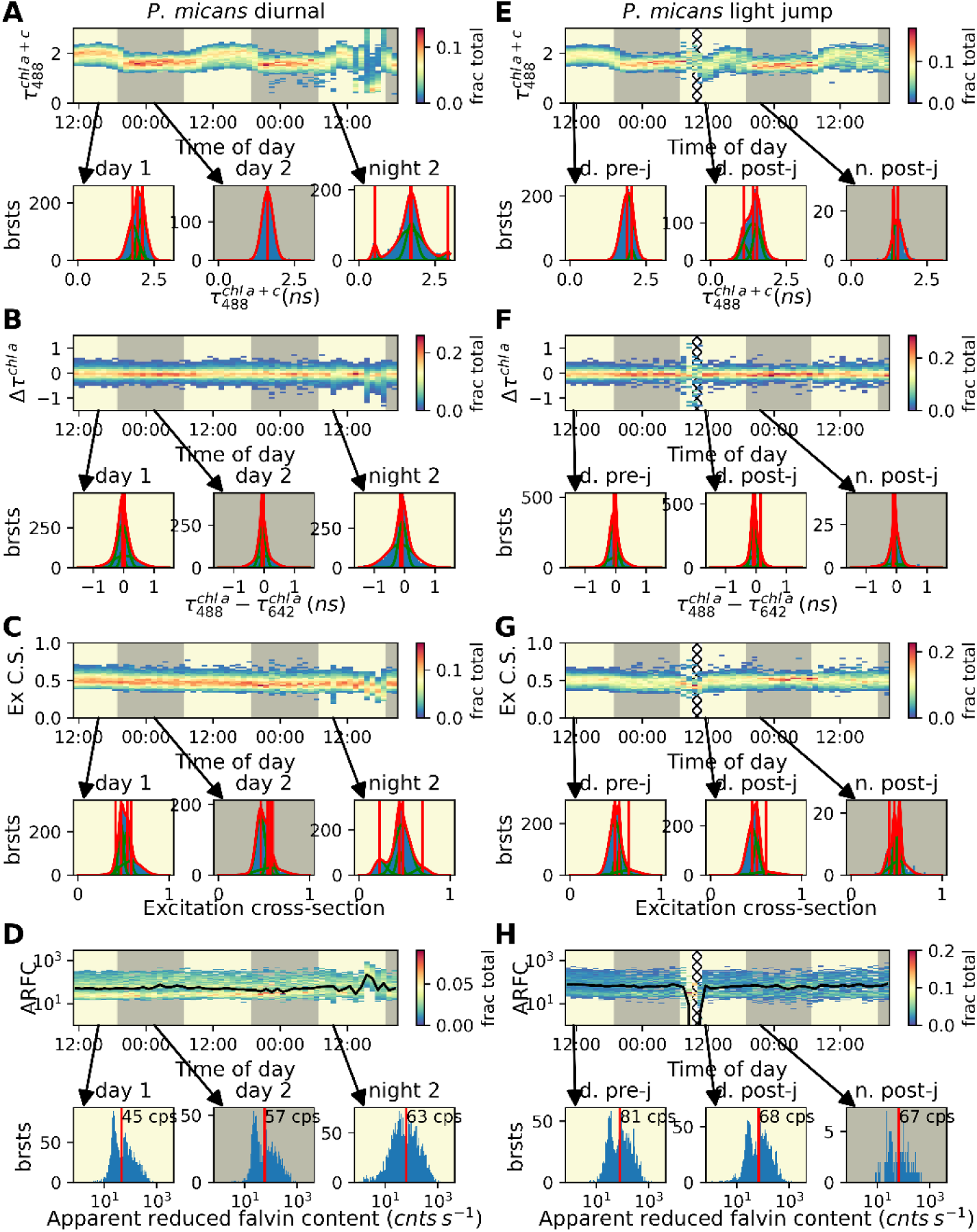
*P. micans* diurnal cycle without or with a light perturbation. (A-D): Diurnal cycle. (E-H): Diurnal cycle in the presence of a light jump perturbation. Kymographs show changes in values of select parameters over time, using hour-long bins. Day and night are indicated by yellow and gray backgrounds respectively, and the period of the light perturbation is indicated by gridded background. Burst histograms of parameters in select hours are shown below each kymograph. Except for D and H, histograms are fit with sum-of-multiple-central-distributions (i.e., specific central distribution selected by parameter) with best-fit selected by F-test. Vertical red lines showing the centroids of each central distribution, and red line showing best-fit model and green lines the underlying central distributions. Parameter selection (for explanation of names see table 2) as follows (A, E) 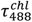, histograms fit with sum-of-multiple-Gaussians, (B, F) 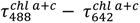, histograms fit with sum-of-multiple-Gaussians, (C, G) Excitation cross-section histograms fit with sum-of-multiple-beta-distributions, and (D, H) apparent reduced flavin content vertical line represents mean value. Towards the end of the diurnal cycle experiment, some clogging has affected the burst rate, and hence the quality of the data.

Lifetime parameters, such as 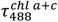, always appeared as a single peak. As for the excitation cross-section, a minor yet nonegligible subpopulation was observed in the histograms throughout the experiment (Fig 7, C, G). While this heterogeneity was less striking relative to that of Syn. WH8102 or *P. purp*, it nevertheless still indicates the presence of cell states.

## Discussion

Our results reveal photo-acclimation strategies in phytoplankton that are characterized by cell-wide coordinated state transitions. Interestingly, we found that PBS-based photosynthetic systems exhibit clearly discernible cell-states, some of which are only populated following an abrupt light intensity perturbation, which induces an increased fluorescence lifetime state. Additionally, the efficiency of the acclimation response following the light perturbation was less than 100%, as can be judged by the fraction of the population of high lifetime cells. The duration of recovery was almost a day. In chlorophyll-based LHC systems, distinct cell states were observed, albeit with smaller differences between cell-state subpopulation values. Upon light perturbation, a unique cell-state with lower lifetime was populated with 100% efficiency, followed by 1-2 hour recovery. Indeed, using the herein presented single-cell approach uncovered novel features of light acclimation responses in different red lineage phytoplankton species.

The emergence of the long LHC fluorescence lifetime subpopulation during the light jump in *Syn.* WH8102 indicates that the acclimation of cyanobacteria to light intensity perturbations involves a choice that occurs independently for each cell. Two possible mechanisms can be considered to explain the reduction in the EET-sensitized emission, combined with the increase in PE fluorescence lifetime values: (1) a reduction of the QY of photochemistry (i.e. accumulation of “closed” photosystems) which, according to eq. 5, would result in longer fluorescence lifetimes, or (2) PE decoupling, in whole or in part from the PBS core (38, 39), which drastically reduces the rate of EET. Considering that cells were dark adapted for 30 minutes in the flow path before detection, the second option should hold true. Closed photosystems revert back to a dark-adapted state within minutes. *Synechoccous* apply a radiative solution to the problem of excess light dissipation. This PE decoupling has to be tied with a signal transduction mechanism that coordinates rapid changes in many photosynthetic units across vast regions of the cell. Recent reports demonstrate the organization of photosynthetic complexes into homogeneous microdomains (40) and suggests that spatio-temporal properties of cell signaling in cyanobacteria can be very robust (41). *P. micans* adapts to light perturbations in a different way, with fluorescence lifetimes decreasing, and negligible EET changes. Therefore, in response to a light jump these dinoflagellates activate NPQ mechanisms(42). Evidence for two distinct sub-populations, based on excitation cross-section, was observed to a certain extent also in the dinoflagellates. The mechanisms described here represent two different acclimation strategies that are relevant to the time scale of changes in light intensity under mixing water column conditions.

All tested organisms showed variability in their parameter values. We note that when taking all parameters together, each organism in each given condition occupies its own value signatures in the parameter space. This makes the approach presented here applicable for environmental studies. Multidimensional fluorescence parameter signatures will identify different genera, experiencing different conditions within complex natural populations. We expect to solve the inverse problem of identifying photosynthetic microorganisms and their cell photophysiological states from multidimensional parameter signatures. While the herein presented setup used two pulsed-interleaved laser excitation sources, four distinct spectral channels and three parameter type bases (i.e., fluorescence intensities, brightnesses and lifetimes), more pulsed-interleaved laser excitation sources and better spectral filtration into more distinct spectral bands, will further increase the information content, which, in turn, will further substantiate the uniqueness of multidimensional parameter signatures. Therefore, the time-resolved multiparameter single cell approach presented here is a basis for functional single cell physiological monitoring as response to a variety of biotic and abiotic perturbations.

## Materials and Methods

### Cell growth

*Synechococcus* WH8102, and *P. purpureum* stock cultures were grown and maintained under blue LED light at 60 μmol photons m^−2^ s^−1^, at 21 °C, with 12\12 light\dark regime. The cultures were shaken continuously, and diluted in artificial seawater medium (ASW) (43) once a week in a sterile glass flask environment.

*Prorocentrum micans* stocks were grown and maintained under blue LED lights at 6 μmol photons m^−2^ s^−1^, at 18 °C, with 12\12 light\dark regime without shaking. f\2 medium (44) was renewed once a week. Cultures were diluted 2-3 days before experiments to ensure the algae are in exponential growth phase and prevent nutrition limitation in the culture.

### Optical setup

A confocal-based setup (ISS™ USA) assembled on top of a modified Olympus IX71 inverted microscope stand, similar to previously reported (45, 46). Excitation was provided by two picosecond pulsed diode lasers (λ = 488 nm, pulse width of 80 ps FWHM, operating at 20 MHz repetition rate, λ = 642 nm, pulse width of 100 ps FWHM, operating at 20 MHz repetition rate QuixX® 488-60 PS and QuixX® 642-140 PS, Omicron-Laserage, GmbH). The laser beams pass through a single-mode polarization maintaining optical fiber (P1-405BPM-FC-Custom, with specifications similar to those of PM-S405-XP, Thorlabs, Newton, NJ, USA) and after passing through a collimating lens (AC080-016-A-ML, Thorlabs), the beam is further shaped by a λ/2 plate (WPMP2-20(OD)-BB 550 nm, Karl Lambrecht Corp., Chicago, IL, USA) and a linear polarizer (DPM-100-VIS, Meadowlark Optics, Frederick, CO, USA). Laser beams are reflected to the optical path through galvo-scanning mirrors (6215H XY, Novanta Corp., Boston, MA, USA) and scan lens (30 mm Dia. x 50 mm FL, VIS-NIR Coated, Achromatic Lens, Edmund Optics, Barrington, NJ, USA; both used for acquiring scanned images in laser scanning mode, LSM), and then into the side port of the microscope body through its tube lens, positioning it at the back aperture of the objective lens using a major dichroic mirror (DM) with high reflectivity at both 488 and 640 nm (ZET488/640m, Chroma, Bellows Falls, Vermont, USA). Laser powers were 130 nW and 300 nW for the 488 nm and 642 nm lasers respectively, measured at the back aperture of the objective. For flow-based experiments, a 20X 0.4 NA air objective lens (PLN20X, Olympus, Japan) was used, while for FLIM measurements a 100X 1.45 NA oil objective lens (UPLSAPO100XO, Olympus, Japan) was used, which focuses the light into a diffraction-limited effective excitation volume, positioned within the sample chamber. A fraction of scattered light returns in the excitation path and is imaged on a CMOS camera (ThorCam, Thorlabs, USA), used for z-positioning, using Airy ring pattern visualization. A fraction of fluorescence from the sample is collected through the same objective lens, is transmitted through the major DM and is focused with an achromatic lens (25mm Dia. x 100 mm FL, VIS-NIR Coated, Edmund Optics) onto a 100 μm diameter pinhole (variable pinhole, motorized, tunable from 20 μm to 1 mm, custom made by ISS^TM^), and then re-collimated with another achromatic lens (f=100 mm; AC254-060-A, Thorlabs, USA). Fluorescence is then chromatically split into four separate detection channels, 510/20, 585/40, 698/70 and 710/40 nm, in the following manner: photons are split using a DM with 652 nm cutoff wavelength (FF652-Di01-25×36, Semrock, USA). Then, photons of the longer wavelength path (>652 nm) are further split in two by a 50:50 beamsplitter (BS; 21000 - 50/50 Beam splitter, Chroma, USA), where photons were selected for longer and shorter wavelengths using bandpass filters 698/70 and 710/40 nm (FF01-710/40-25 and FF01-698/70-25, Semrock, USA) before the detectors. Photons of the shorter wavelength path (<652 nm) are split by a DM with a cutoff wavelength of 555 nm (FF555-Di03-25×36, Semrock, USA), followed by bandpass filters 510/20 nm for photons with wavelength <555 nm, and 585/40 nm for photons with wavelength >555 nm and <652 nm (FF03-510/20-25 and FF01-585/40-25, Semrock, USA) before the detectors. Photons of the four different detection channels are detected using four cooled hybrid photomultipliers (Model R10467U-40, Hamamatsu, Japan), routed through a 4-to-1 router to a time-correlated single photon counting (TCSPC) module (SPC-150, Becker & Hickl, GmbH) as its START signal (the STOP signal is routed from the pulsed laser controller). We perform data acquisition using the VistaVision software (version 4.2.095, 64-bit, ISS^TM^, USA) in the time-tagged time-resolved (TTTR) file format. After data acquisition, data files are converted into the photon HDF5 format (47) for easy dissemination of raw data. Burst analysis was performed using FRETbursts (48).

For fluorescence lifetime imaging (FLIM), Images were attained by using a laser scanning module (LSM), in which a 3-axis DAC module (custom made by ISS^TM^, USA) synchronized the data acquisition and control over the x and y galvo-scanning mirrors, which assisted in bringing the effective excitation volume to different positions to acquire pixel data per a given z layer. Fluorescence lifetime images were attained by taking the mean nanotime of photons in each photon stream (combination of excitation laser and detection channel) per pixel, taking pixels with a threshold of 200 photons. Mean nanotimes were used in leu of the more common tail-fitting in order to unify parameter calculation methods between FLIM and flow based measurements. Images were acquired with 0.1 ms pixel dwell times in 30 x 30 μm^2^ image dimensions and 256 x 256 pixels (6.55 s per frame), summing 30 such frame scans per image (196.6 s per image). Excitation powers and filter setup were identical to the ones used in flow-based acquisition.

### Calculation of burst fluorescence parameters

Acquired data was first converted into photon HDF5 format (47) using *phconvert*. Converted files were then processed with Jupyter notebooks using the FRETBursts python package (48) to perform burst search. All channel burst search was performed with sliding window of size m=20 and with the minimum criterion of an instantaneous photon counting rate of F=6 times the background rate. Finally, bursts were selected (i.e., gated) such that they had >50 photons in the 488 nm excitation 565-605 nm emission and the 488 nm excitation 665-733 nm emission streams, and >1 photon in streams with 665-733 nm or 690-730 nm emission, and that background correction did not result in the 488 nm excitation 565-605 nm emission stream having a negative value.

The mean fluorescence lifetime parameters are calculated based on the mean of all photon nanotimes (i.e., the mean photon nanotime) of all the photons longer than a given threshold set by the IRF, in a burst. If the background rate is low, relative to the fluorescence rate in bursts, then the effect of the few background photons in each burst on the quantification of the mean photon nanotime becomes negligible. In such conditions, it can be shown that the value of the mean photon nanotime equals the mean fluorescence lifetime, even if the underlying fluorescence decay exhibits multiple exponential components. The mean fluorescence lifetime parameters are calculated per each photon stream (i.e., each combination of excitation source and detection channel) and are named accordingly (see Table 1).

The number of photons in a burst, known as the burst size, and the time from the first detected photon of a burst to the last photon of the burst, known as the burst duration, are calculated. Then, the mean brightness of the burst is the ratio of the burst size and its duration. The brightness ratio parameters are calculated as ratio of burst brightness values for different photon streams and are named accordingly (see Table 2).

### Model fitting to histograms of burst fluorescence parameter values

Each histogram of a given burst fluorescence parameter in this work is not only presented but also undergoes model fitting to a model of a sum-of-multiple-central-distributions function (Gaussian or Beta). Model fitting is conducted for from one to four central distributions, and then the F-test is applied to select the best-fit function. When plotting, we include a red vertical line to indicate position of each central distribution, the value of which we refer to as the centroid in figure legends. For Gaussian distributions the mean and mode are the same, and thus we choose this value as the centroid. For beta distributions, the mode is the more visually-prominent value, and thus we choose to represent the centroid as the mode. Thus, the centroid is always the mode. For the apparent reduced flavin content, we do not perform model fitting, and thus take the mean of the raw data to generate a representative value for the data.

Gaussian functions were chosen to describe mean nanotime data, owning to the central limit theorem, while for brightness ratio parameters, which are by definition limited to a range of 0 - 1, a beta functions were used. It should be noted that, in some cases, the best-fit functions do not perfectly describe the underlying histogram data. While we do report the results of these model fittings, we majorly report the parameter value histograms themselves, and rely mostly on them for interpretations, and the best-fit Gaussian means are shown as guides to the eye.

### Flow-based Data Acquisition

To acquire fluorescence photon bursts of cells in flow-based experiments, an environment chamber was prepared comprising of a box within which blue LED mats (49) were secured at the top to provide appropriate light intensities for the diurnal cycle in each experiment. The power supply was linked to a timer which activated the light at 7:00 am (7:00) each day and deactivated at 7:00 pm (19:00) each day. A computer fan provided air circulation within the environment chamber.

A cell culture flask was modified with an outlet at the bottom, to which a 1 mm inner diameter FPLC tube was attached. The cell culture flask was placed on a magnetic stir plate and a small oblate spheroid stir bar was added to agitate the flask at 100 rpm, the slowest speed the stir plate could provide. The FLPC tube (0.5 mm inner diameter PEEK tubing) was passed through a small hole made in the box and linked to a microfluidic channel (80666, Ibidi). Silicone tubing (1.6 mm inner diameter; 10842, Ibidi) was used to connect the outlet of the microfluidic channel to a syringe pump which was operated with a 10 mL syringe at a flow rate of 7 μL min^−1^, corresponding to horizontal velocity of 1.12 mm s^−1^. All connections to the microfluidic chamber were made through Luer adapters.

The stage was manually aligned to position the focal volume near the beginning of the channel in the direction of flow, and laterally in the middle of the channel. Similarly, the focal position was chosen to be in the middle between the top and bottom of the channel. This ensures that single cell bursts are from a region devoid of surface effects and reducing the time between cells leaving the flask and their measurement.

### Bulk spectroscopy

Fluorescence excitation and emission spectra of the cultures was measured with PTI Quantamaster spectrometer (Horiba, Kyoto Japan), using glass cuvettes. Using a peristaltic pump, culture was fed from a culture flask into the cuvette through silicone tubing (1.6 mm inner diameter; 10842, Ibidi). Conditions such as temperature, illumination and stirring were adjusted to match the conditions of the confocal single cell experiments. Three measurements were taken every 30 min for a period of 48 hours. Excitation spectra were measured at 700 nm emission detection. Emission spectra were measured using 480 nm excitation, as well as using 642 nm excitation. Slit widths were fixed to 5 nm.

### Conditions for Cell Imaging

To acquire images of cyanobacteria, we adapted the procedure developed by Iermak and coworkers(50). Cells are spatially confined in agarose inside custom chambers. Chambers were assembled by first punching a hole through a 0.5 mm silicone sheet (JTR-S-0.5, Grace Biolabs, USA), which was then placed over an 18 x 18 mm^2^ coverslip. The hole in the silicone sheet provided a chamber which was partially filled with 8 μL of ASW agarose, and then another coverslip was placed on top, and pressed to seal the chamber until the agarose set solid. *Syn.* WH8102 cells are concentrated by centrifuging 1 mL of culture volume at 16,000 rcf for 3 minutes. Then, 990 μL of supernatant was removed and the pellet resuspended in the remaining 10 μL of solution. Then the top coverslip was pried off to expose a flat agar surface onto which 2 μL of the concentrated culture was deposited. The top coverslip is then reapplied, and the full assembly secured in a holder that both serves as a mount to position the chamber on the microscope and provide pressure to compress the chamber together and spatially confine the cells.

## Data, Materials, and Software Availability

Code and data for this study are archived at DOI https://doi.org/10.5281/zenodo.10614780

## Author Contributions

Conceptualization: PDH, NK, EL; Methodology: PDH, NBE; Investigation: PDH, NBE, NK, EL; Visualization: PDH, NBE; Funding acquisition: PDH, NK, EL; Project administration: NK, EL; Supervision: NK, EL; Writing – original draft: PDH, NK, EL; Writing – review & editing: PDH, NBE, NK, EL

## Classification

Biological Sciences, Plant Biology

## Acknowledgments

This work was supported by the Israel Science Foundation (grant no. 663/23 to NK, no. 556/22 to EL), the Racah Nano Venture Fund (to NK and EL) and the Zuckerman Stem Leadership Postdoctoral Program (to PDH).

