## Supplementary material for "Multiparameter-based photosynthetic state transitions of single phytoplankton cells": SI

#### Supplementary Information

### Supplementary Figures

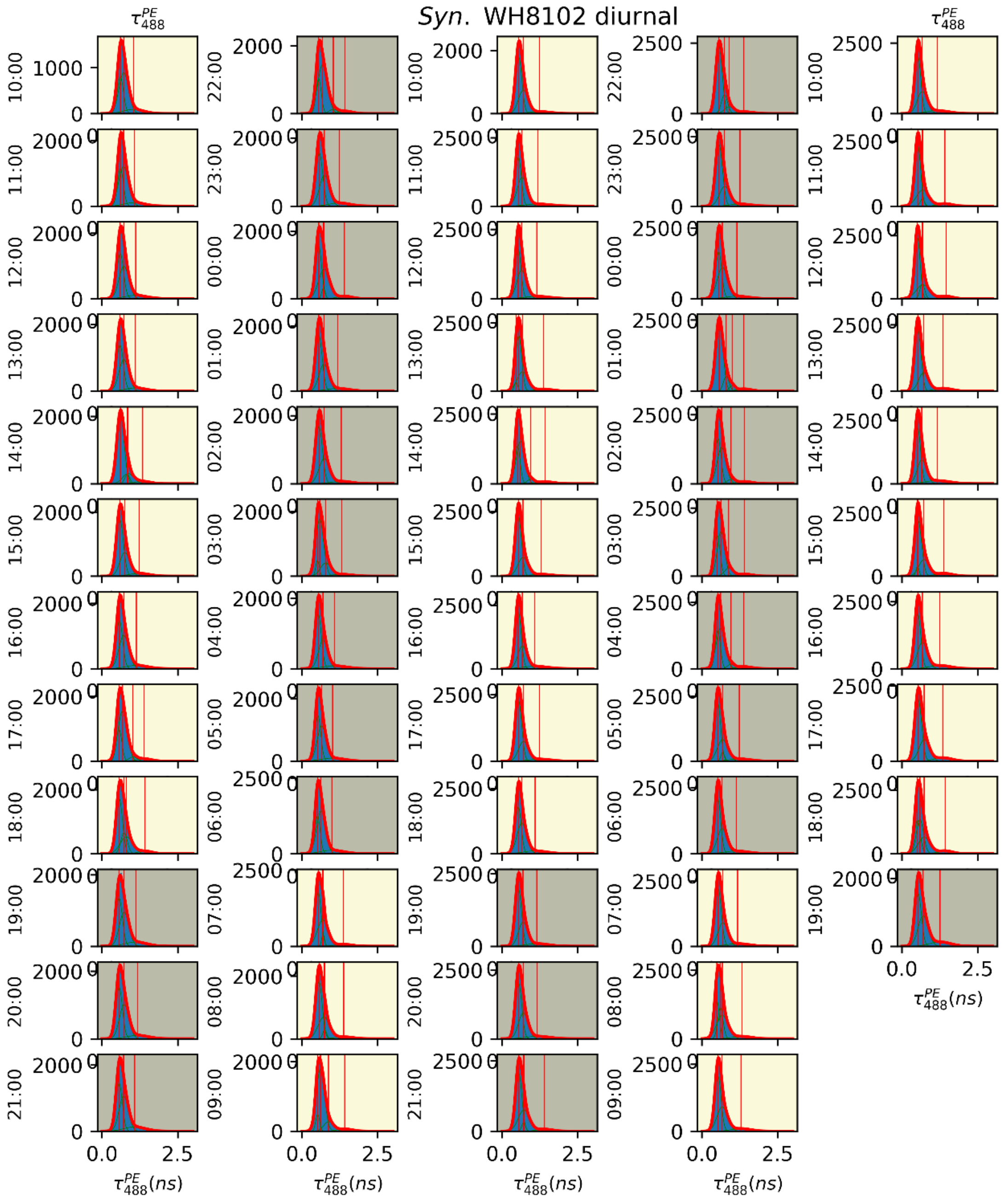


**Fig. S1.** Histograms of mean fluorescence lifetime $\tau_{488}^{PE}$ during diurnal cycle of *Syn*. WH8102 (supplement to Fig 3). Day/night and model fittings as in Fig. 3. Red line is best-fit as determined by F-test of sum-of-multiple-Gaussians-models, green lines are underlying Gaussians composing best-fit, and vertical red lines are the centroids of the underlying Gaussians.

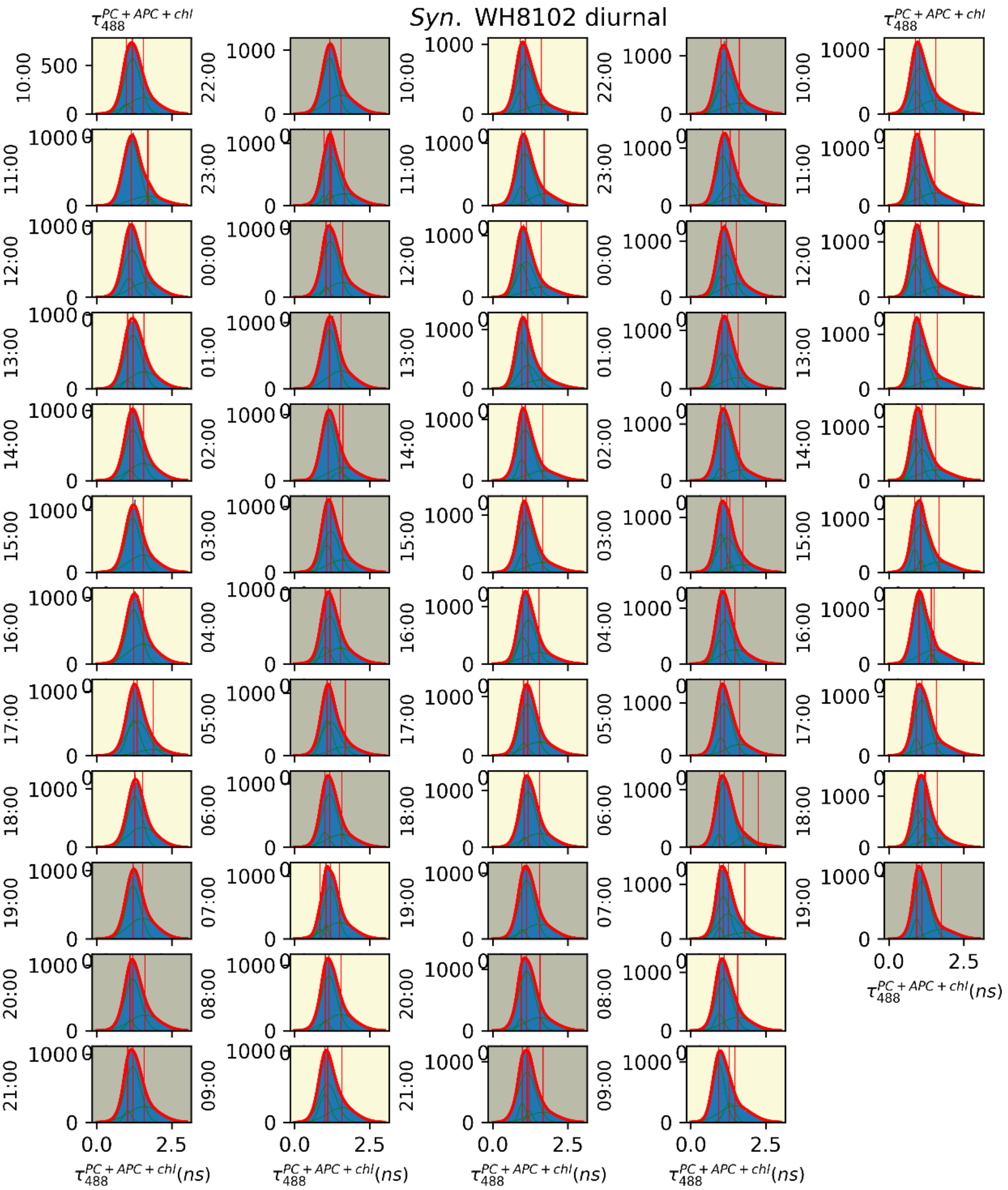


**Fig. S2.** Histograms of mean fluorescence lifetime $\tau_{488}^{PC+APC+chl}$ during diurnal cycle of *Syn.* WH8102 (supplement to Fig. 3). Day/night and model fittings as in Fig. 3. Red line is best-fit as determined by F-test of sum-of-multiple-Gaussians-models, green lines are underlying Gaussians composing best-fit, and vertical red lines are the centroids of the underlying Gaussians.

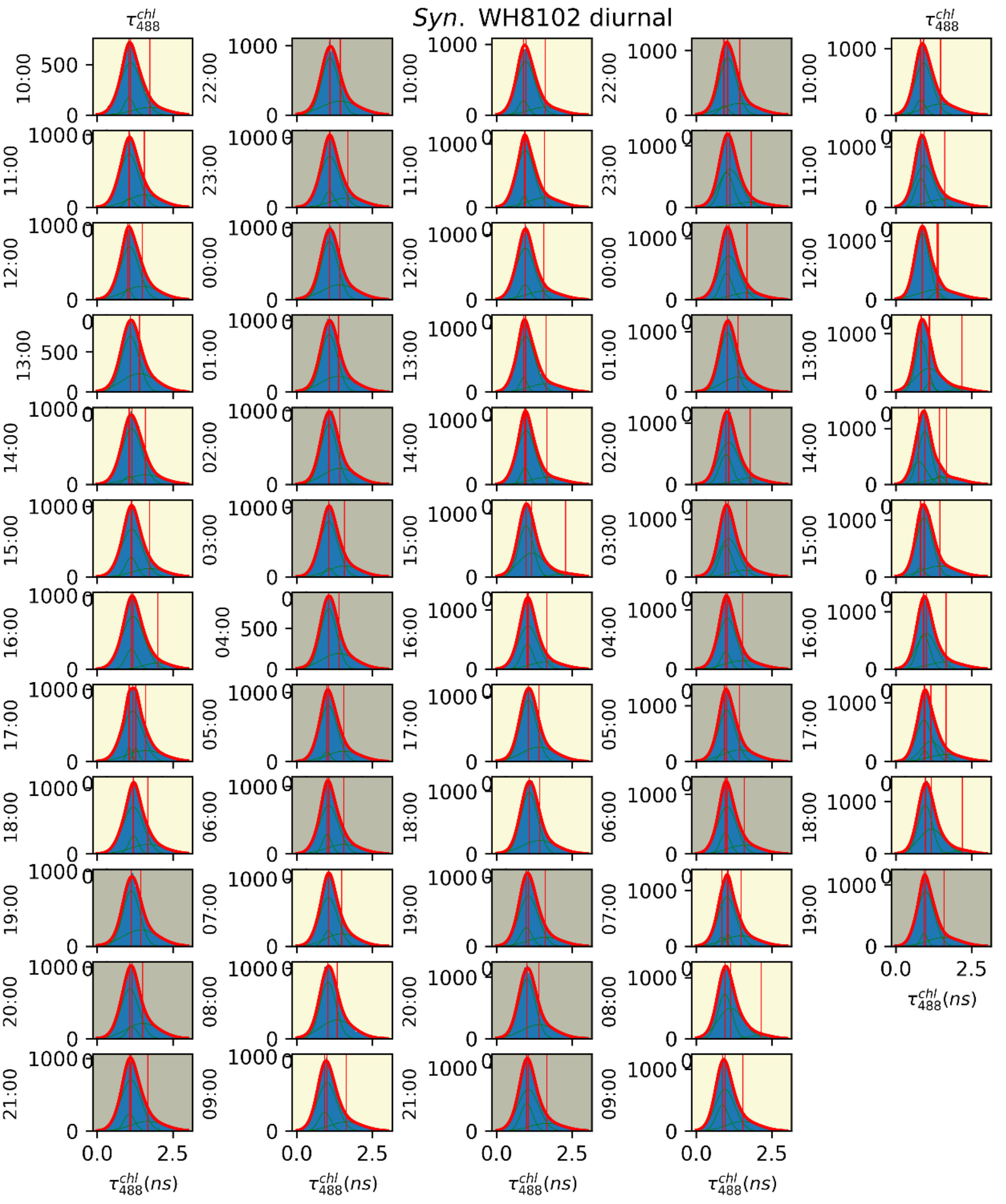


**Fig. S3.** Histograms of mean fluorescence lifetime $\tau_{488}^{chl}$ during diurnal cycle of *Syn.* WH8102 (supplement to Fig. 3). Day/night and model fittings as in Fig. 3. Red line is best-fit as determined by F-test of sum-of-multiple-Gaussians-models, green lines are underlying Gaussians composing best-fit, and vertical red lines are the centroids of the underlying Gaussians.

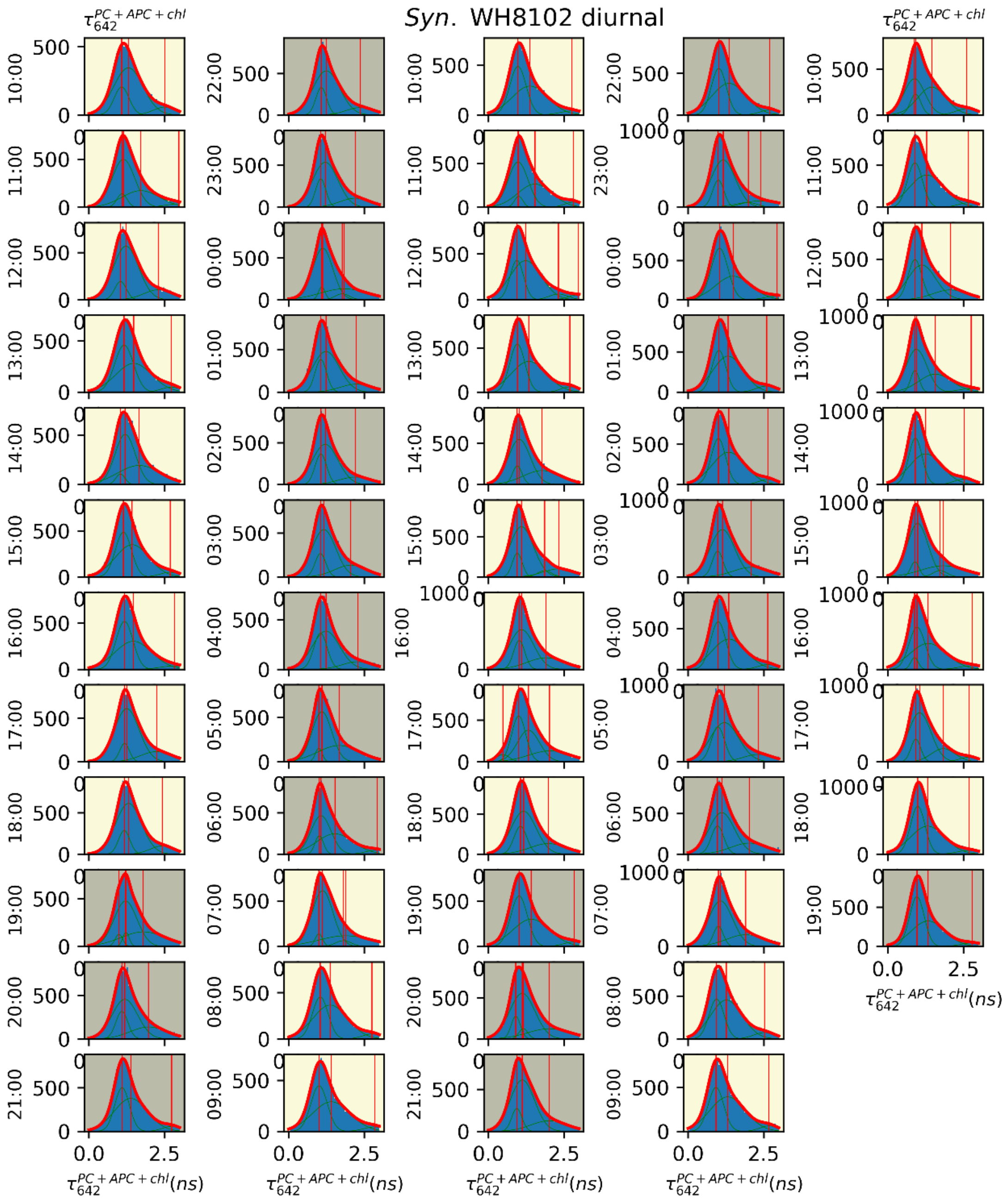


**Fig. S4.** Histograms of mean fluorescence lifetime $\tau_{642}^{PC+APC+chl}$ during diurnal cycle of *Syn.* WH8102 (supplement to Fig. 3). Day/night and model fittings as in Fig. 3. Red line is best-fit as determined by F-test of sum-of-multiple-Gaussians-models, green lines are underlying Gaussians composing best-fit, and vertical red lines are the centroids of the underlying Gaussians.

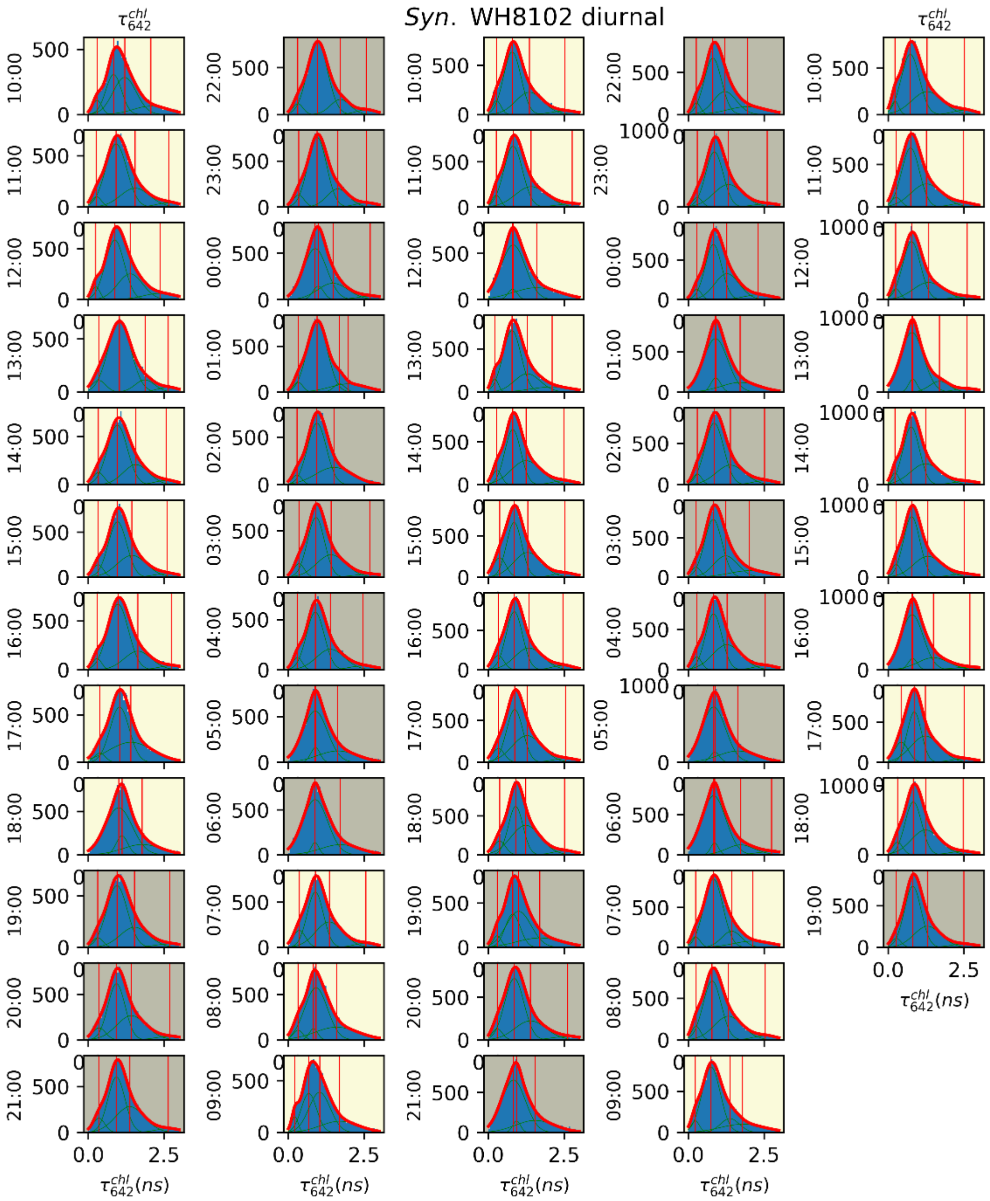


**Fig. S5.** Histograms of mean fluorescence lifetime $\tau_{642}^{chl}$ during diurnal cycle of *Syn.* WH8102 (supplement to Fig. 3). Day/night and model fittings as in Fig. 3. Red line is best-fit as determined by F-test of sum-of-multiple-Gaussians-models, green lines are underlying Gaussians composing best-fit, and vertical red lines are the centroids of the underlying Gaussians.

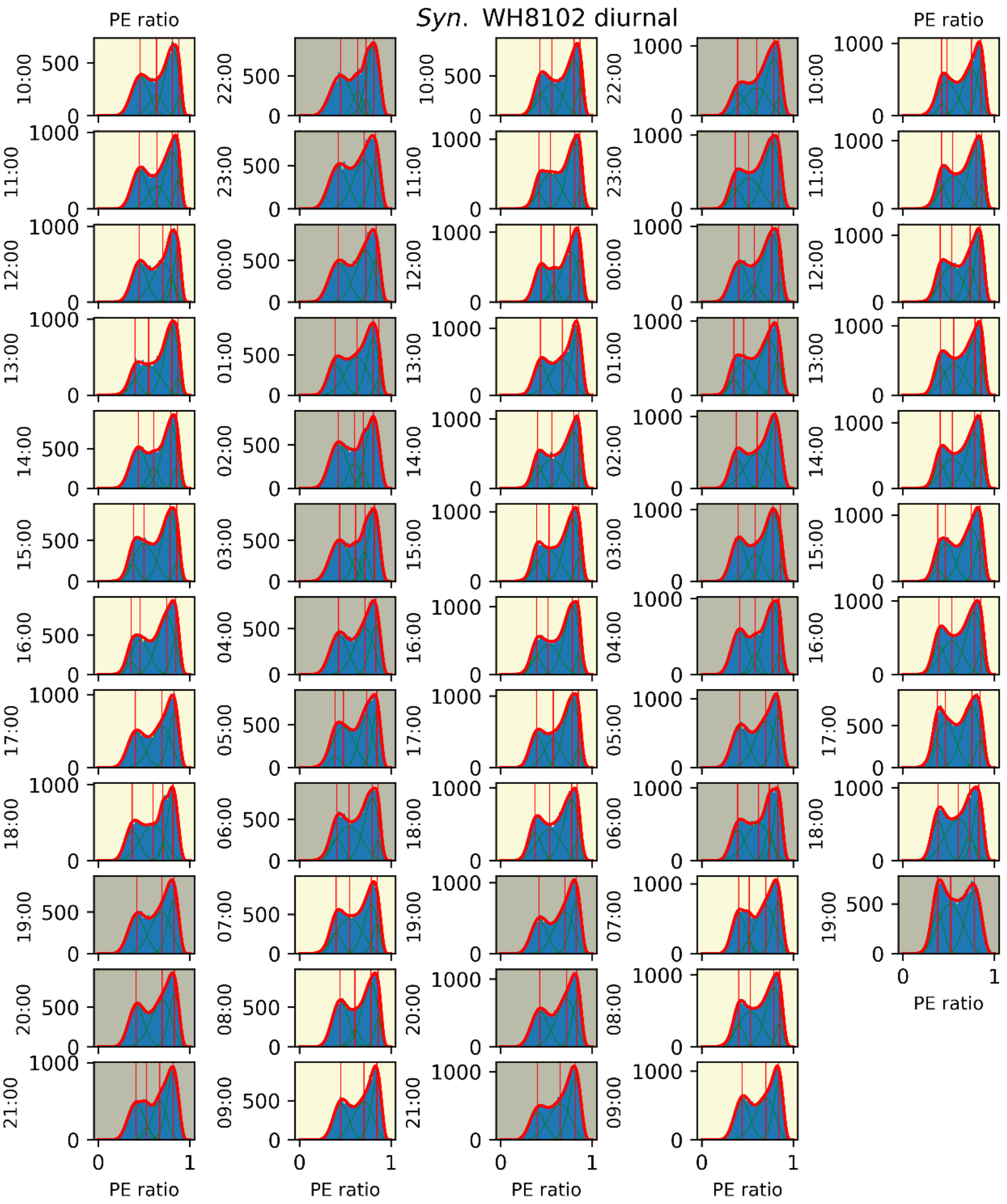


**Fig. S6.** Histograms of PE ratio during diurnal cycle of *Syn*. WH8102 (supplement to Fig. 4). Day/night and model fittings as in Fig. 4. Red line is best-fit as determined by F-test of sum-of-multiple-beta-distribution-models, green lines are underlying beta distributions composing best-fit, and vertical red lines are the centroids of the underlying beta distribution.

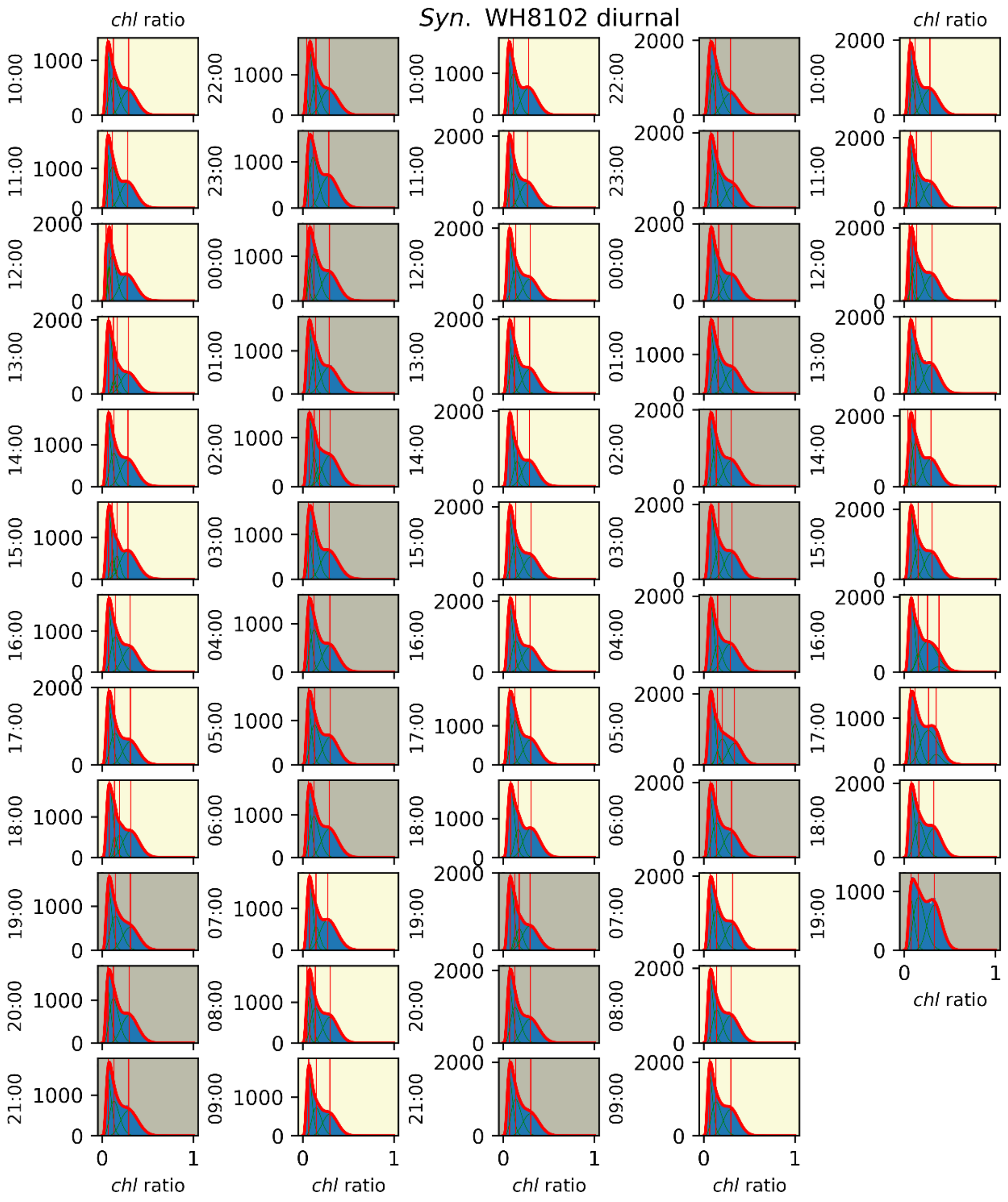


**Fig. S7.** Histograms of *chl* ratio during diurnal cycle of *Syn*. WH8102 (supplement to Fig. 4). Day/night and model fittings as in Fig. 4. Red line is best-fit as determined by F-test of sum-of-multiple-beta-distribution-models, green lines are underlying beta distributions composing best-fit, and vertical red lines are the centroids of the underlying beta distribution

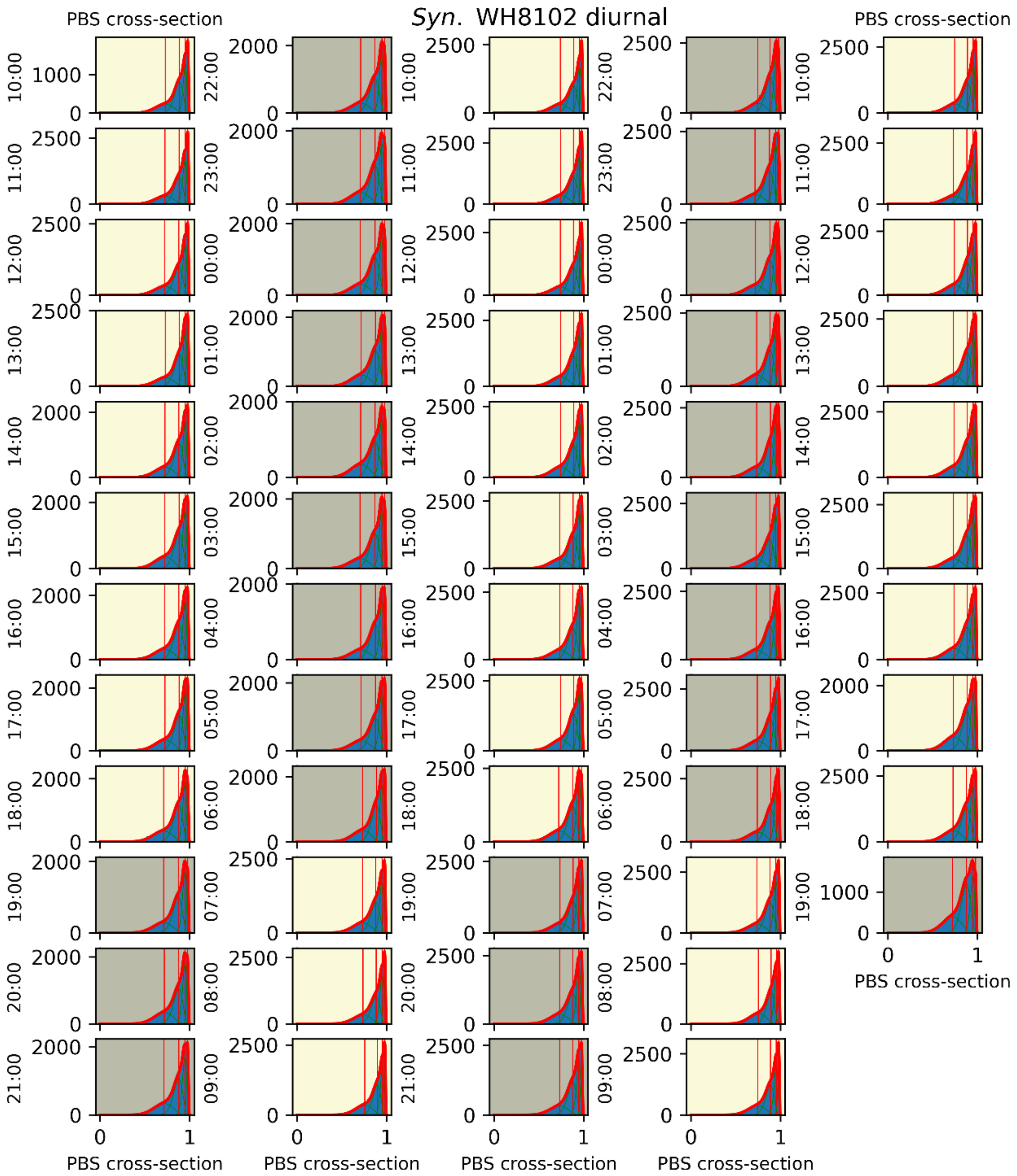


**Fig. S8.** Histograms of PBS cross-section during diurnal cycle of *Syn.* WH8102 (supplement to Fig. 4). Day/night and model fittings as in Fig. 4. Red line is best-fit as determined by F-test of sum-of-multiple-beta-distribution-models, green lines are underlying beta distributions composing best-fit, and vertical red lines are the centroids of the underlying beta distribution

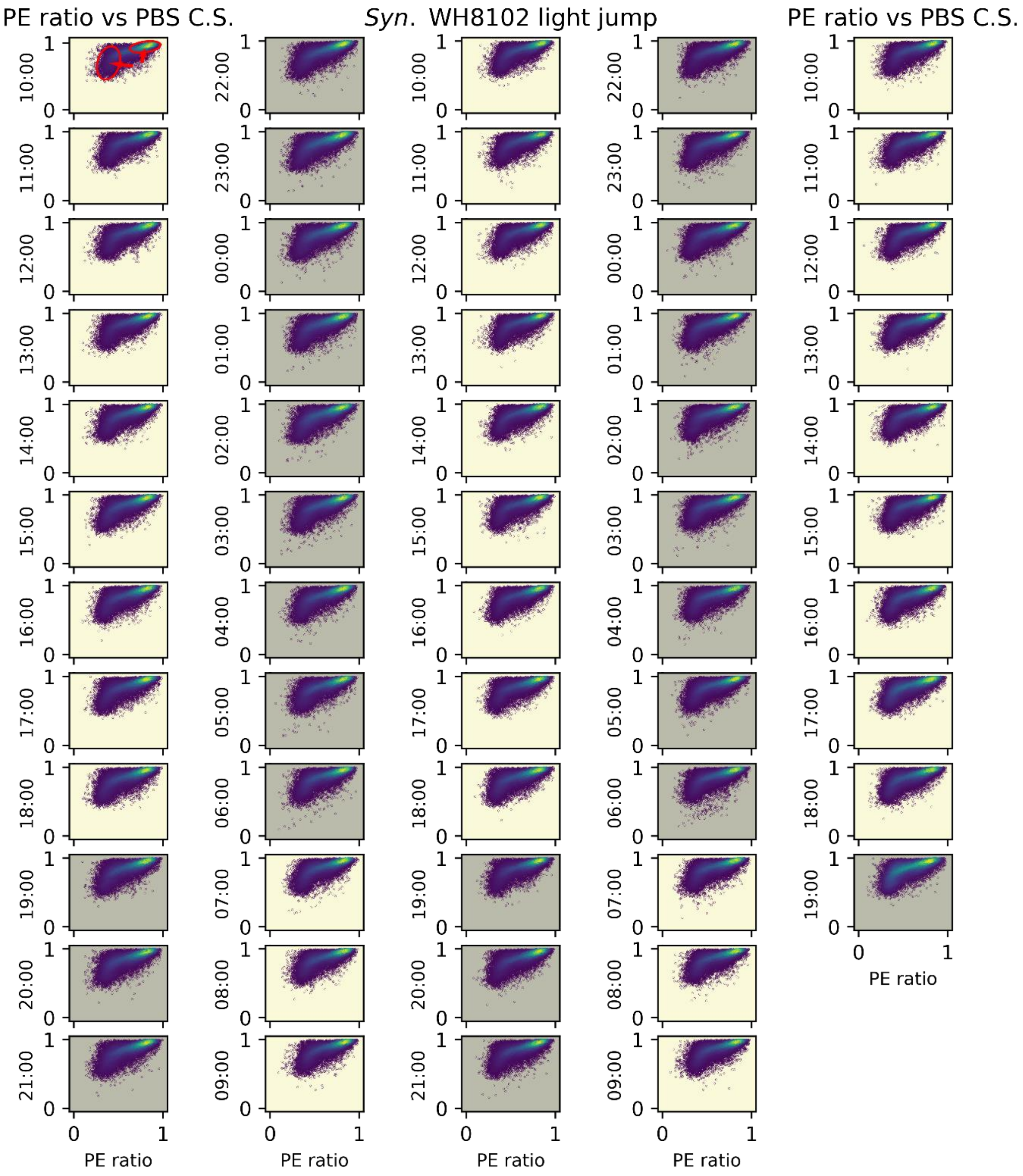


**Fig. S9.** Scatter plots of PE ratio:PBS cross-section during diurnal cycle of *Syn.* WH8102. Yellow background denotes day, gray background denotes night. In upper left plot circles and ellipses indicate suggested subpopulations of cells. The ellipse in the upper right contains 50% of the bursts in the plot, while the center left ellipse contains 23% of the bursts

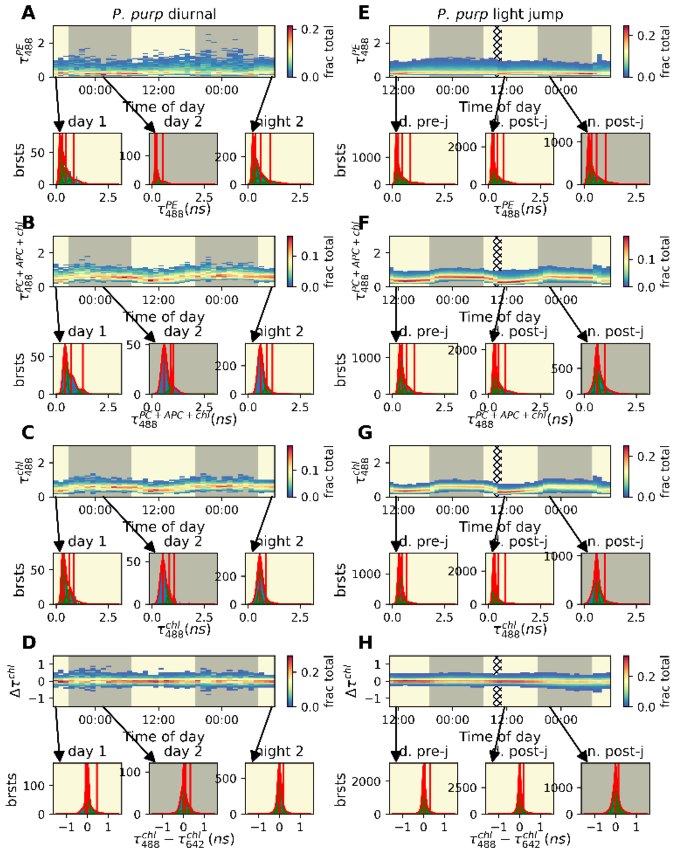

**Fig. S10.** Kymographs selected lifetime and lifetime diference parameters of *P. purp* during (A-D) diurnal cycle and (E-H) light jump perturbation. Display/fitting procedures follows same rules as figs 2-5,7. Selected parameters: (A, E) $\tau_{488}^{PE}$, (B,F), (C,G) PBS cross-section, and (D, H) apparent reduced flavin content. Towards the end of the diurnal cycle experiment after light jump, some clogging has affected the burst rate, and hence the quality of the data. Histograms of selected 1 hour time periods displayed below kymorgraphs, for A-C and E-G (brightness ratio parameters) fittings to sum-of-multiple-beta-distribution models, with best-fit selected by F-test are displayed as red lines, with underlying beta-distributions plotted as green lines, and the centroids marked with vertical red lines.

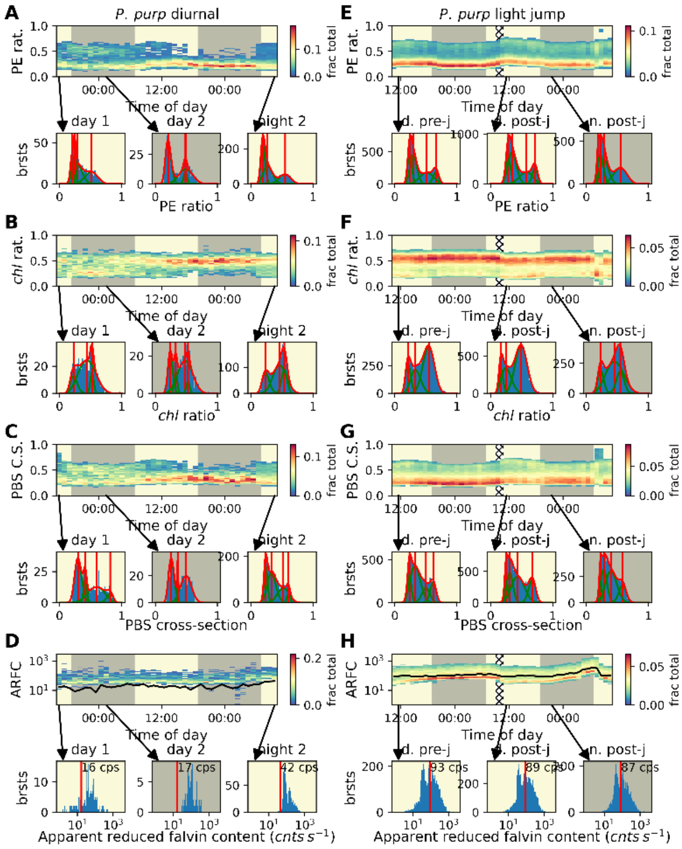


**Fig. S11.** Kymographs selected brightness ratio and brightness parameters of *P. purp* during (A-D) diurnal cycle and (E-H) light jump perturbation. Display/fitting procedures follows same rules as figs 2-5,7. Selected parameters: (A, E) PE ratio, (B, F) *chl* ratio, (C, G) PBS cross-section, and (D, H) apparent reduced flavin content. Towards the end of the diurnal cycle experiment after light jump, some clogging has affected the burst rate, and hence the quality of the data. Histograms of selected 1 hour time periods displayed below kymorgraphs, for A-C and E-G (brightness ratio parameters) fittings to sum-of-multiple-beta-distribution models, with best-fit selected by F-test are displayed as red lines, with underlying beta-distributions plotted as green lines, and the centroids marked with vertical red lines.

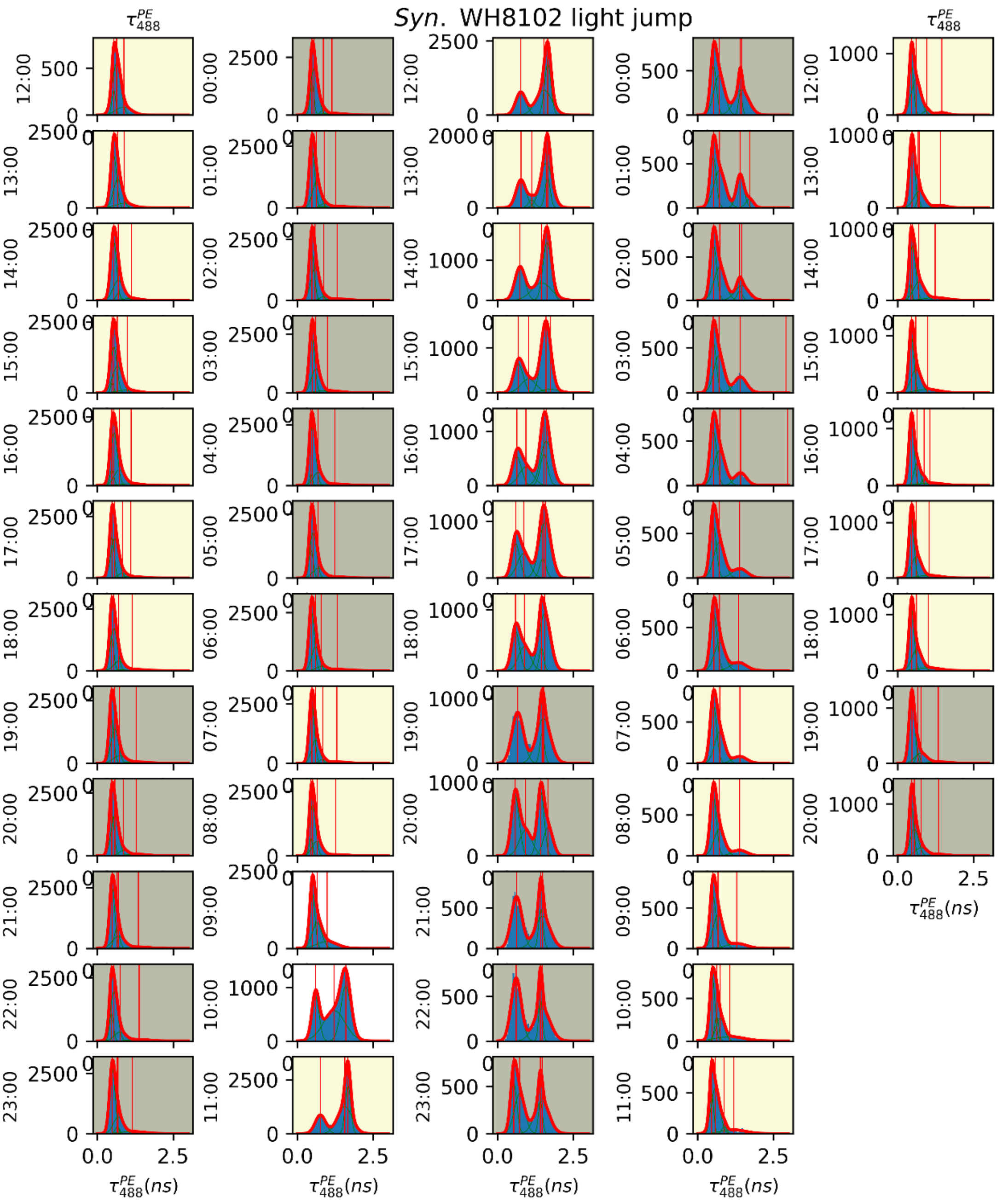


**Fig. S12.** Histograms of mean fluorescence lifetime $\tau_{488}^{PE}$ during diurnal cycle of *Syn*. WH8102, before, during and after a light jump perturbation (supplement to Fig. 5). Day/night/light jump and model fittings as in Fig. 5. Red line is best-fit as determined by F-test of sum-of-multiple-Gaussians-models, green lines are underlying Gaussians composing best-fit, and vertical red lines are the centroids of the underlying Gaussians.

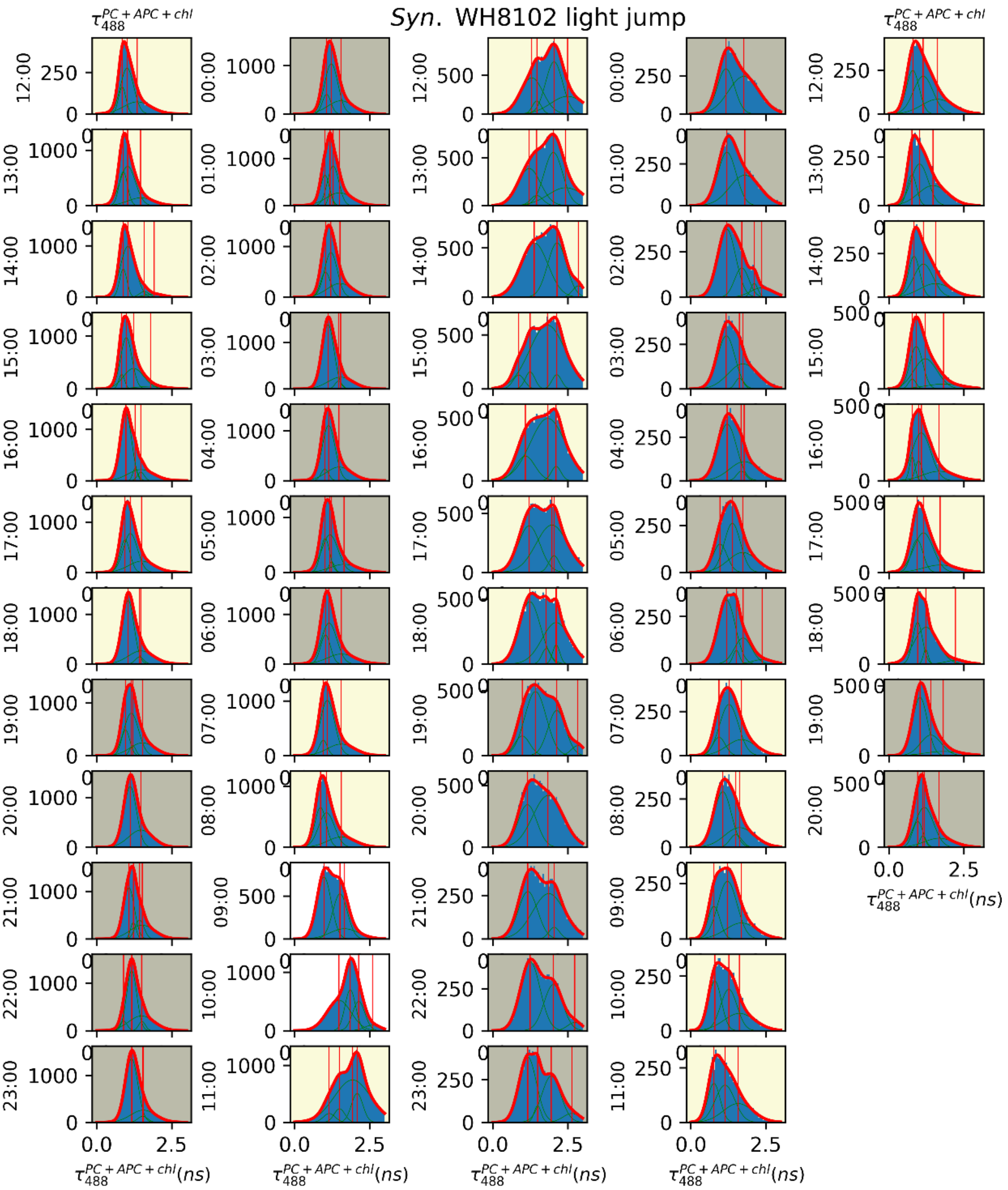


**Fig. S13.** Histograms of mean fluorescence lifetime $\tau_{488}^{PC+APC+chl}$ during diurnal cycle of *Syn.* WH8102, before, during and after a light jump perturbation (supplement to Fig. 5). Day/night/light jump and model fittings as in Fig. 5. Red line is best-fit as determined by F-test of sum-of-multiple-Gaussians-models, green lines are underlying Gaussians composing best-fit, and vertical red lines are the centroids of the underlying Gaussians.

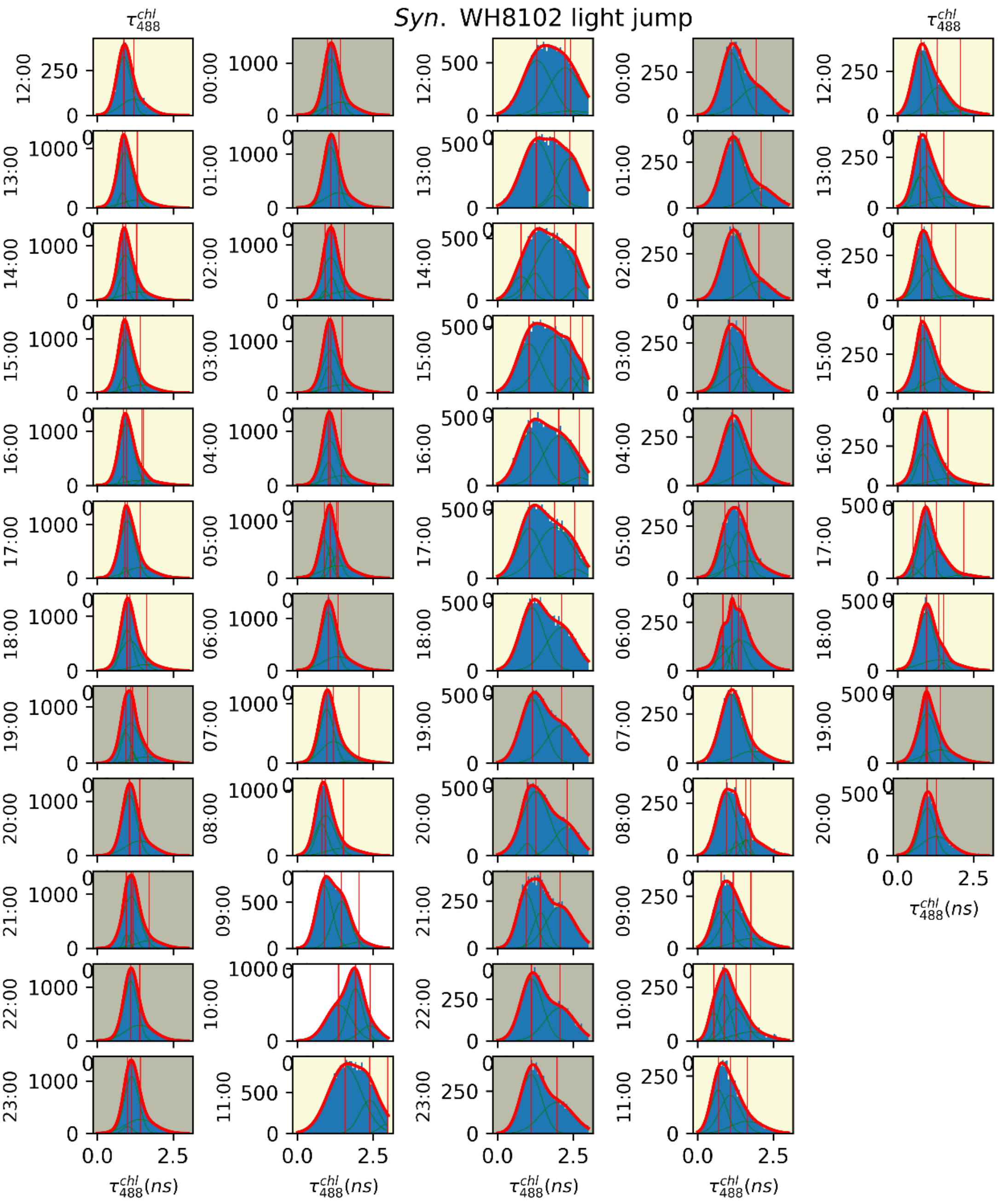


**Fig. S14.** Histograms of mean fluorescence lifetime $\tau_{488}^{chl}$ during diurnal cycle of *Syn*. WH8102, before, during and after a light jump perturbation (supplement to Fig. 5). Day/night/light jump and model fittings as in Fig. 5. Red line is best-fit as determined by F-test of sum-of-multiple-Gaussians-models, green lines are underlying Gaussians composing best-fit, and vertical red lines are the centroids of the underlying Gaussians.

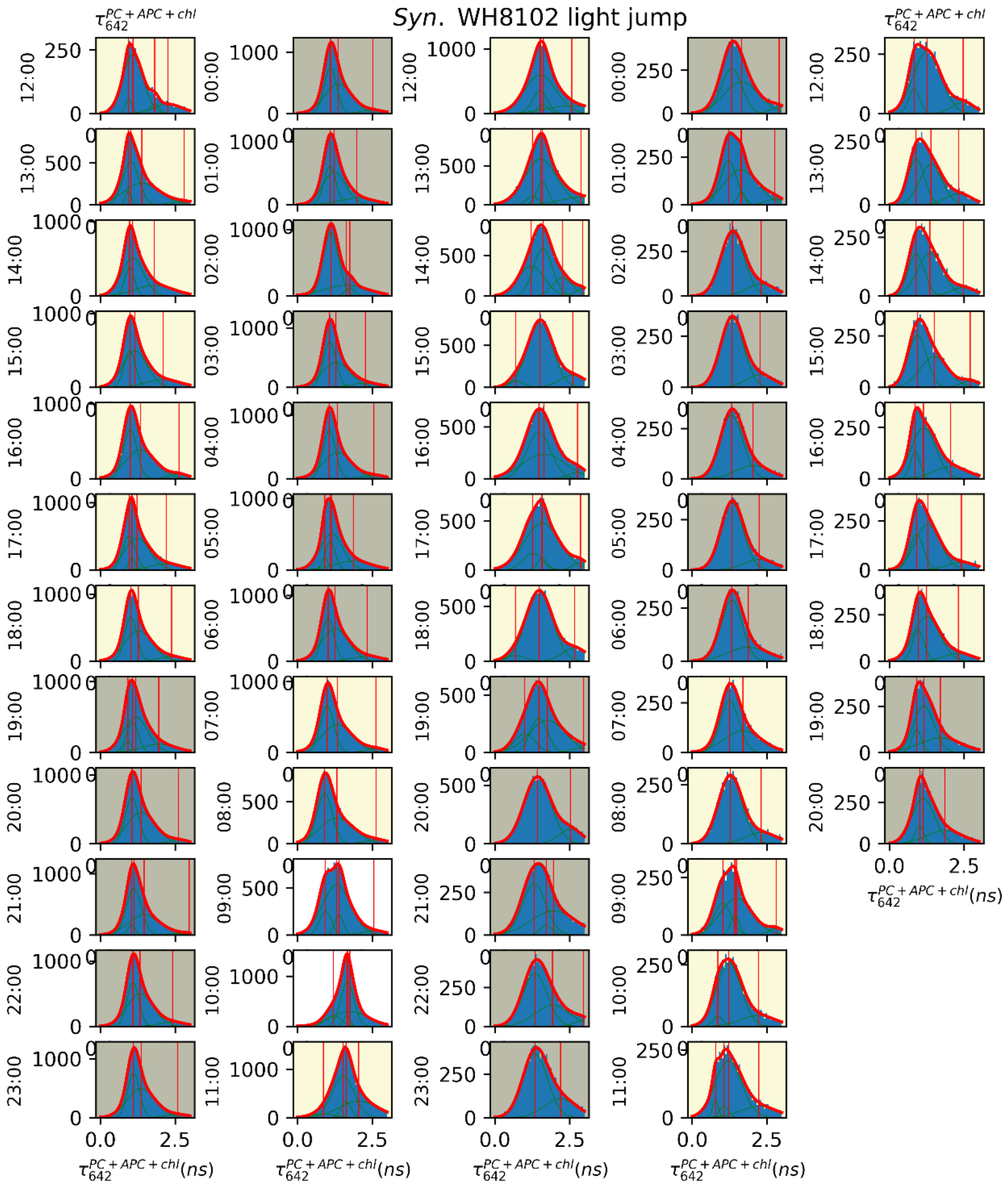


**Fig. S15.** Histograms of mean fluorescence lifetime $\tau_{642}^{PC+APC+chl}$ during diurnal cycle of *Syn.* WH8102, before, during and after a light jump perturbation (supplement to Fig. 5). Day/night/light jump and model fittings as in Fig. 5. Red line is best-fit as determined by F-test of sum-of-multiple-Gaussians-models, green lines are underlying Gaussians composing best-fit, and vertical red lines are the centroids of the underlying Gaussians.

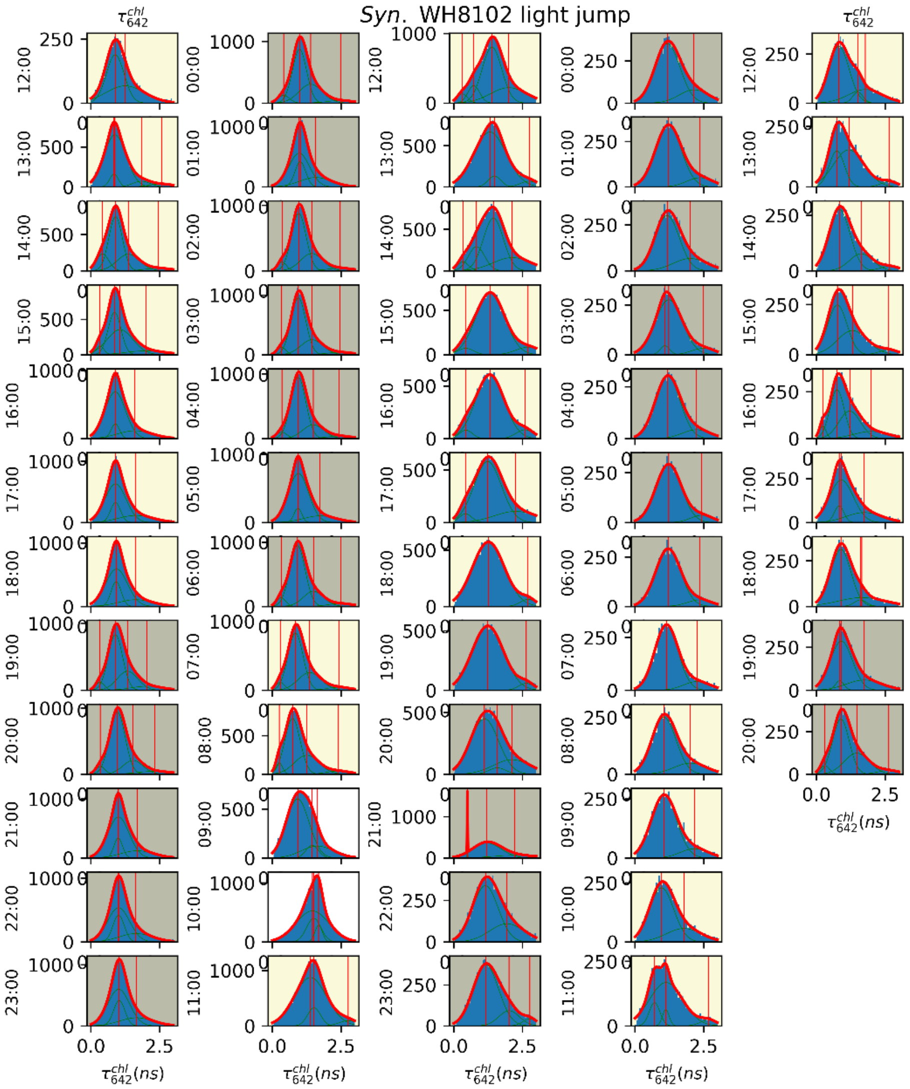


**Fig. S16.** Histograms of mean fluorescence lifetime $\tau_{642}^{chl}$ during diurnal cycle of *Syn*. WH8102, before, during and after a light jump perturbation (supplement to Fig. 5). Day/night/light jump and model fittings as in Fig. 5. Red line is best-fit as determined by F-test of sum-of-multiple-Gaussians-models, green lines are underlying Gaussians composing best-fit, and vertical red lines are the centroids of the underlying Gaussians.

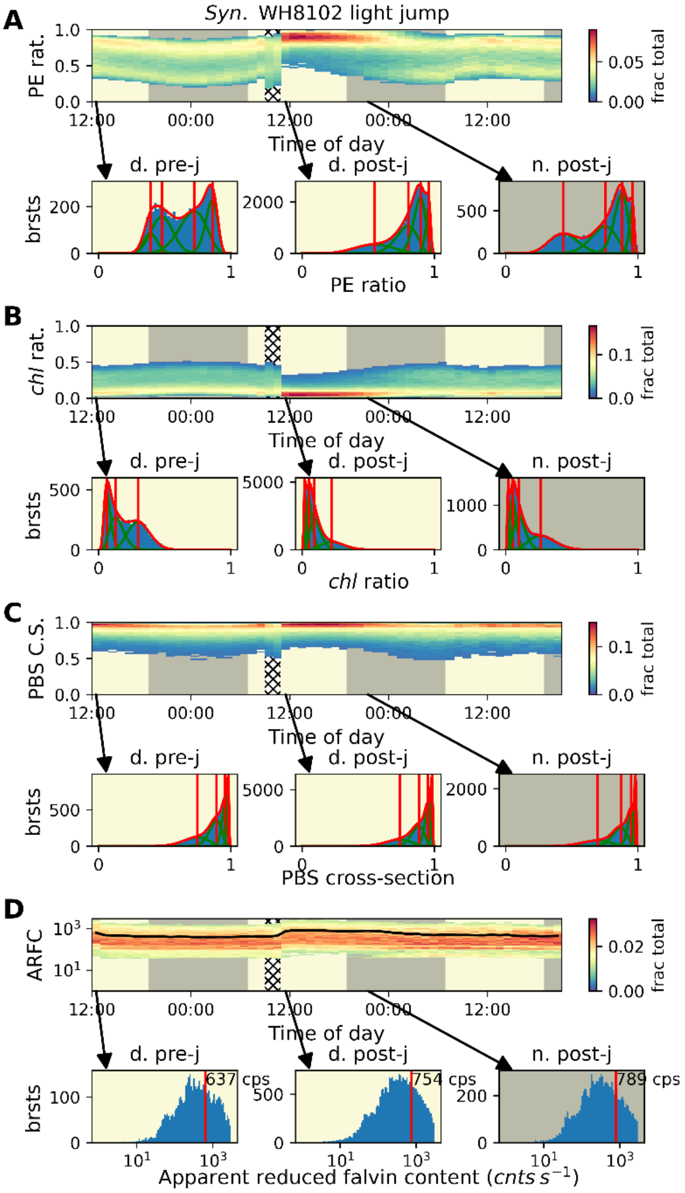


**Fig. S17.** Kymographs of brightness ratio and brightness parameters of during diurnal cycle of *Syn*. WH8102, before, during and after a light jump perturbation, parameter organization identical to Fig. 4, light jump is indicated by gridded background. Histograms of bursts below kymographs show select hour periods of brightness ratio parameters with fittings to models of a sum-of-multiple-beta-distributions, with the best-fit model selected by the F-test. Vertical lines in histograms represent the centroids beta distributions when performing a sum-of-multiple-beta-distributions model fitting to the data and selecting the best-fit model based on F-test, red line represents the fitted model, while green lines represent the beta distributions within each model. The (A) PE ratio (i.e., relative proportion of emission from PE), (B) *chl* ratio (i.e., relative emission from *chl a*), (C) PBS cross-section (i.e., relative fluorescence from 488 nm excitation), and (D) apparent reduced flavin content (i.e., brightness of emission in flavin channel), are shown. In histograms of apparent reduced flavin content the mean of raw data is reported as a vertical red line, instead of fitting the day to any type of distribution.

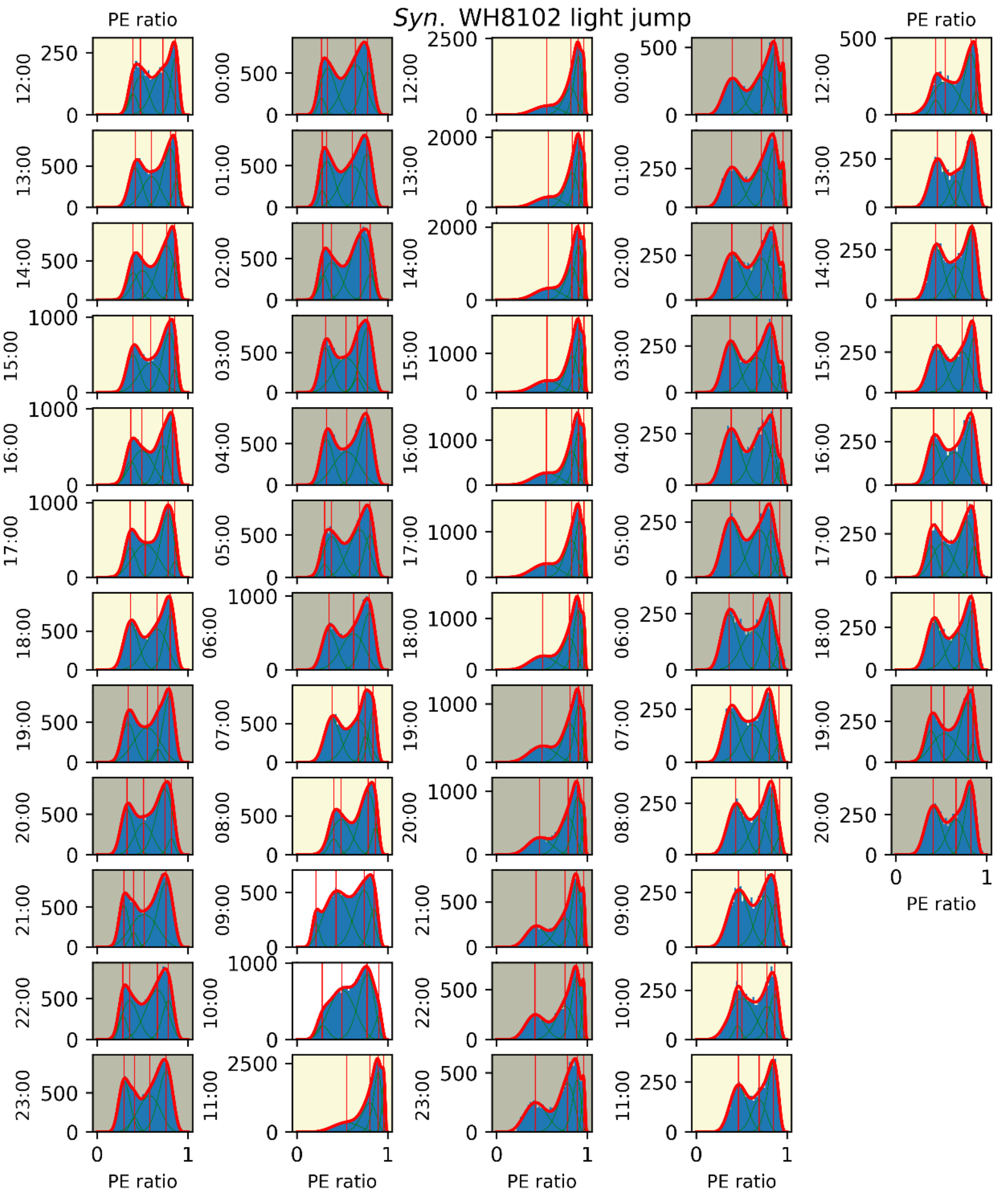


**Fig. S18.** Histograms of PE ratio during diurnal cycle of *Syn*. WH8102, before, during and after a light jump perturbation. Day/night and model fittings as in Fig. 4. Red line is best-fit as determined by F-test of sum-of-multiple-beta-distribution-models, green lines are underlying beta distributions composing best-fit, and vertical red lines are the centroids of the underlying beta distribution.

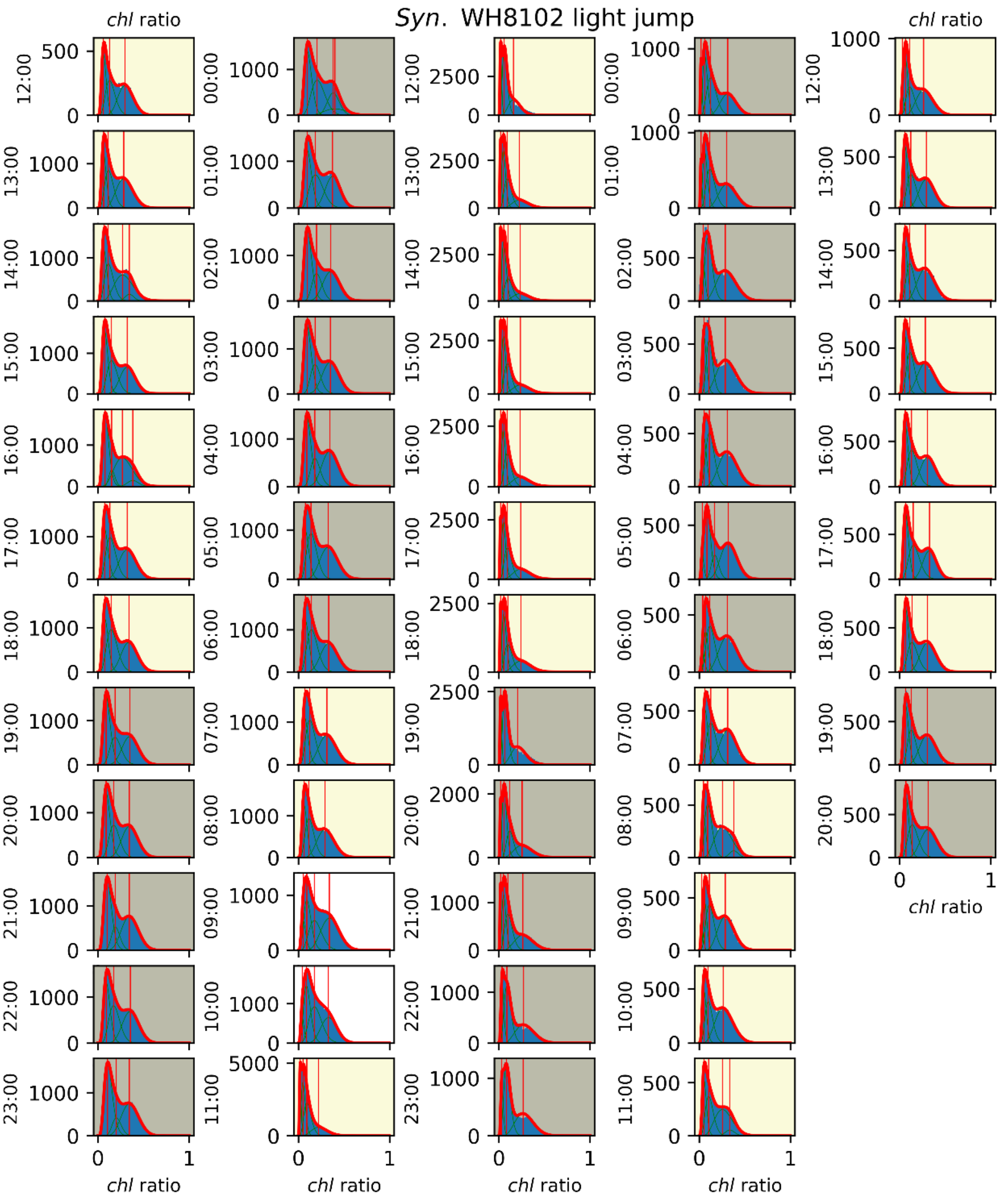


**Fig. S19.** Histograms of *chl* ratio during diurnal cycle of *Syn*. WH8102, before, during and after a light jump perturbation. Day/night and model fittings as in Fig. 4. Red line is best-fit as determined by F-test of sum-of-multiple-beta-distribution-models, green lines are underlying beta distributions composing best-fit, and vertical red lines are the centroids of the underlying beta distribution.

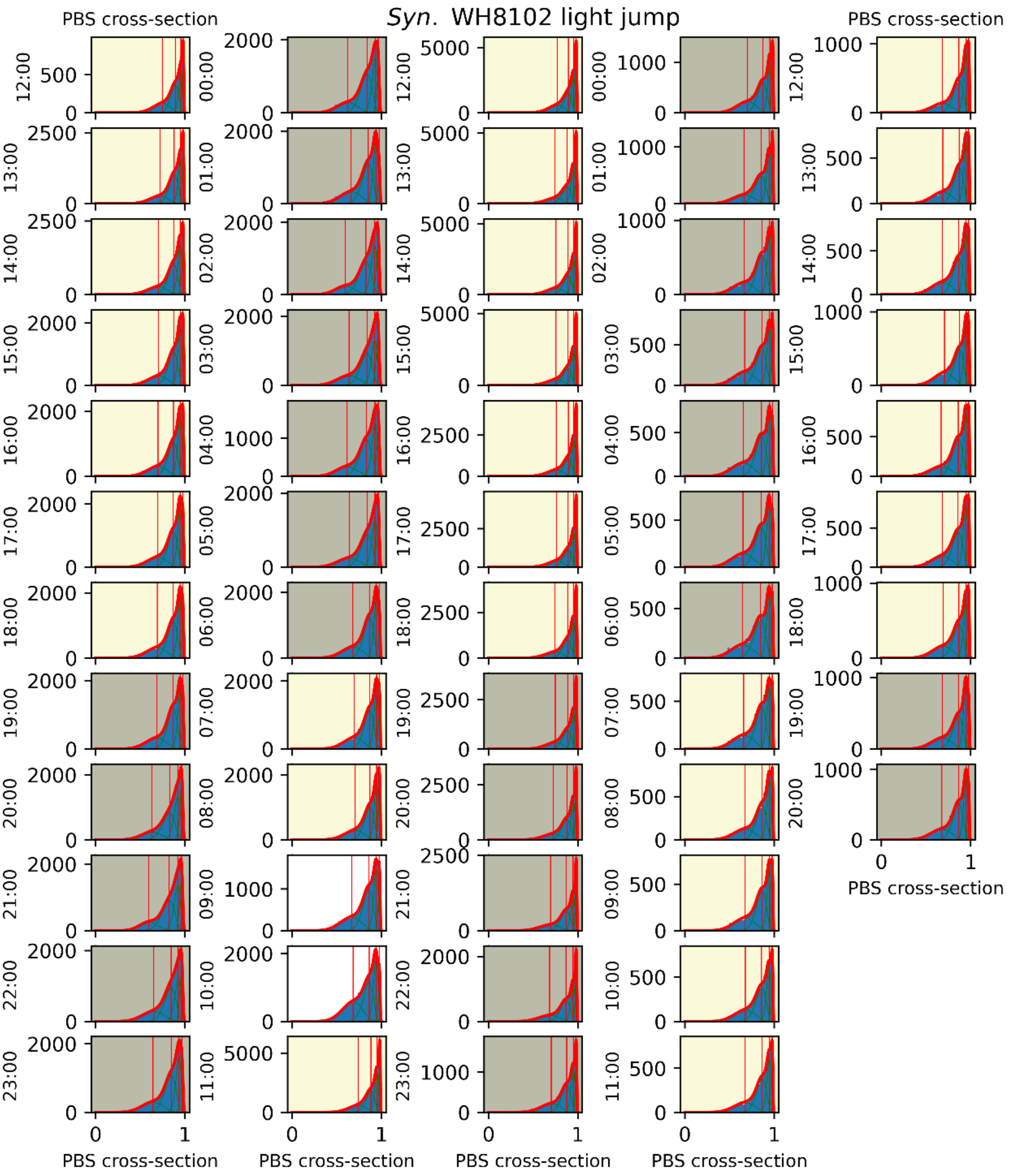


**Fig. S20.** Histograms PBS cross-section during diurnal cycle of *Syn*. WH8102, before, during and after a light jump perturbation. Day/night and model fittings as in Fig. 4. Red line is best-fit as determined by F-test of sum-of-multiple-beta-distribution-models, green lines are underlying beta distributions composing best-fit, and vertical red lines are the centroids of the underlying beta distribution.

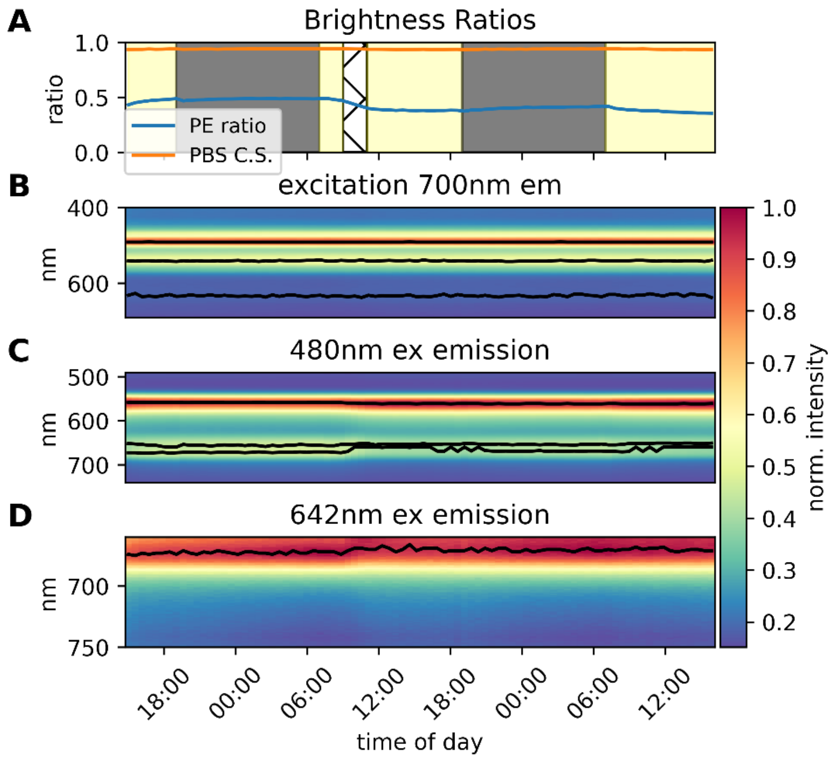


**Fig. S21.** ***Syn.*** **WH8102 bulk excitation spectra measured every 30 minutes.**
(A) Brightness ratios equivalent to those measured in single organism measurements, brightnesses calculated as integrals over windows identical to filters used in single organism measurements, yellow background indicates day, gray indicates night and gridded background indicates light jump, (B) excitation spectra, at $\lambda_{em.}$=700 nm, (C) emission spectra, at $\lambda_{ex.}$=480 nm, (D) emission spectra, at $\lambda_{ex.}$=642 nm. Black lines indicate local maxima of spectra throughout the day.

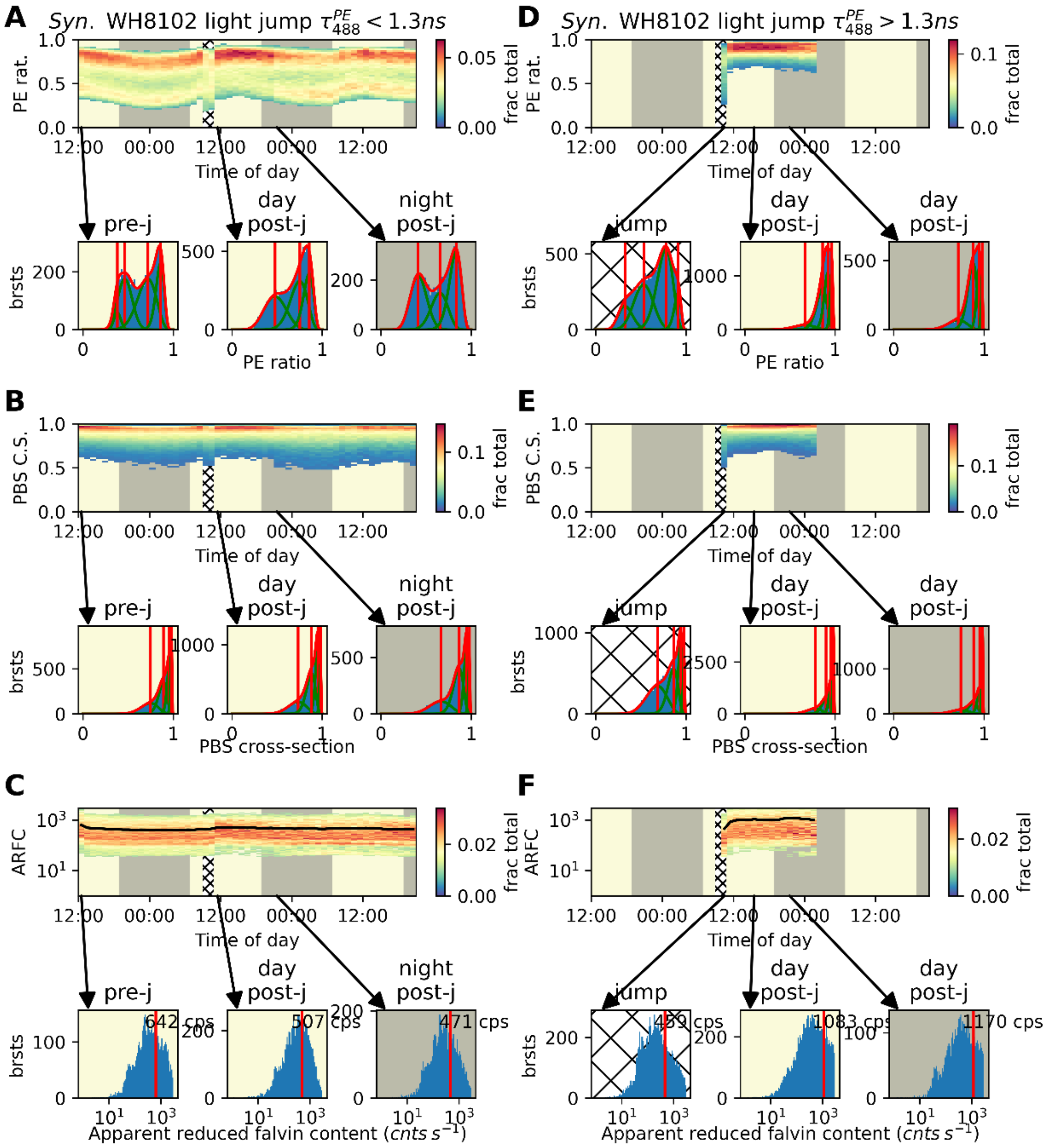


**Fig. S22.** Behavior of long (D-F) and short (A-C) mean fluorescent lifetime $\tau_{488}^{PE}$ subpopulations of *Syn.* WH8102 in light jump experiment. Different parameters shown in rows: PE ratio (A, D), PBS cross-section (B, E), apparent reduced flavin content (C, F). (A, C) organisms with $\tau_{488}^{PE} < 1.3\ ns$, (B, D) $\tau_{488}^{PE} > 1.3\ ns$, data only displayed if >2,000 bursts are present in a given population in a given hour. Histograms below kymograph show selected hour-long periods within the experiment. For PE ratio and PBS cross-section, histograms are fit to sum-of-multiple-Gaussian or -beta-distributions respectively, with best-fit selected using F-test. Vertical red lines indicate centroids of each central distribution in model, red line indicates best fit and green lines the underlying central distributions. For apparent reduced flavin content, the mean value of raw data is reported instead.

### Supplementary Tables

| ***Table S1: Information on chromophores and pigments contributing to fluorescence spectra*** | | | |
|---|---|---|---|
| Chromophore/pigment | Photosynthetic complexes | ~$\lambda_{max.\ abs.}$ (nm)[1] | ~$\lambda_{max.\ fluo.}$ (nm)[2] |
| *Phycoerythrobilin (PEB) and Phycourobilin (PUB)* | *Phycoerythrin (PE)* | *495, 540* | *560-570* |
| *Phycocyanobilin (PCB)* | *phycocyanin (PC)* | *565, 620* | *650* |
| *Phycocyanobilin (PCB)* | *allophycocyanin (APC)* | *640* | *670* |
| *Chlorophyll a (chl a)* | *Photosystem II (PSII), peridinin-chlorophyll a-binding protein (PCP), chlorophyll a-chlorophyll c2-peridinin-protein (acpPC).* | *430, 660* | *670 -680* |
| *Chlorophyll c (chl c)* | *chlorophyll a-chlorophyll c2-peridinin-protein (acpPC).* | *450, 630* | *650* |

*The phycobillisomes (PBS) of marine Synechococcus and red algae are attached to the thylakoid membrane surface and covalently bind open tetrapyrrole chromophores. Dinoflagellates utilize Chlorophylls a and c (chl a and c) and the carotenoid peridinin in their two LHCs, PCP and acpPC. It is noteworthy that Photosystem I (PSI) do not contribute significant fluorescence at room temperature. Photosystem chlorophyll a (chl a) emission is dominated by PSII.*

[1] Approx. absorption wavelength maxima
[2] Approx. fluorescence wavelength maxima

**Table S2.** Day/night mean fluorescence lifetime parameters of *Syn.* WH8102 diurnal cycle.

| | **day** | | | **night** | | | **Welch's T-test** | |
|---|---|---|---|---|---|---|---|---|
| | **mean** | **std err** | **# bursts** | **mean** | **std err** | **# bursts** | **statistic** | **P-value** |
| $\tau_{488}^{PE}$ | 0.669 | $2.9\times10^{-5}$ | $7.2\times10^{7}$ | 0.669 | $2.1\times10^{-5}$ | $1.4\times10^{8}$ | -0.767 | 0.443 |
| $\tau_{488}^{PC+APC+chl}$ | 1.265 | $5.2\times10^{-5}$ | $7.2\times10^{7}$ | 1.265 | $3.7\times10^{-5}$ | $1.4\times10^{8}$ | -2.389 | 0.017 |
| $\tau_{488}^{chl}$ | 1.168 | $5.8\times10^{-5}$ | $7.2\times10^{7}$ | 1.169 | $4.1\times10^{-5}$ | $1.4\times10^{8}$ | -2.318 | 0.020 |
| $\tau_{642}^{PC+APC+chl}$ | 1.431 | $9.8\times10^{-5}$ | $7.2\times10^{7}$ | 1.431 | $7.0\times10^{-5}$ | $1.4\times10^{8}$ | 0.140 | 0.888 |
| $\tau_{642}^{chl}$ | 1.181 | $1.0\times10^{-4}$ | $7.2\times10^{7}$ | 1.181 | $7.1\times10^{-5}$ | $1.4\times10^{8}$ | -0.495 | 0.620 |

Mean is over all bursts detected in every day/night. Welch's T-test was performed on these values, and the large number of bursts accounts for the unusually small P-values given the nearly similar means.
